# Lift&Add - rapid and robust addition of new species to alignments of conserved non-coding sequences

**DOI:** 10.1101/2025.10.14.682260

**Authors:** Navya Shukla, Irene Gallego Romero

## Abstract

Identifying sequence constraint across long evolutionary distances is a powerful method for the discovery of functional genomic sequences, especially putative non-coding elements. Conserved elements have been a mainstay of comparative genomic research, and can be further investigated for specific-specific sequence acceleration to dissect the genetic basis of trait evolution. The conclusions of these comparative genomic studies are however contingent in on the number and range of species included in this phylogenetic analysis. One group of species that has been largely under-represented in genomic comparisons are the marsupials, due to the dearth of marsupial genomes in most publicly available whole-genome alignments. In this study, we firstly showed how biased phylogenetic distributions can profoundly affects estimations of conservation/acceleration with a focus on the marsupials. Then we present a bioinformatic workflow that rapidly enabled us to map 13,287 vertebrate conserved elements—a majority of which were intergenic—identified from the 60-species whole-genome alignment of vertebrates (containing only 4 marsupials) to up to 12 new marsupial genomes ("Lift"). Following this, we combined these new marsupials sequences back to multiple species alignments of these conserved elements ("Add"). Lastly, we demonstrate with our test dataset how expanding phylogenetic breadth can change the conclusions of a comparative genomic analysis.

## Introduction

The growing availability of genome-scale data in the last two decades has transformed both the breadth and resolution of comparative genetic investigations. Genome alignments of more than two species can clarify patterns of molecular evolution and have been particularly instrumental in the analysis of non-coding elements, by allowing the identification of intergenic elements that have undergone little sequence divergence over long evolutionary distances [1–7]. These conserved non-coding elements are presumed to be of functional importance and can be further examined for lineage-specific variation—an excess of substitutions, also known as sequence acceleration. Acceleration can be indicative of positive selection, and has been used to isolate non-coding elements potentially underlying human-specific traits [8, 9], wing development in bats [10] and hibernation in independent lineages of placental mammals [11].

Over the last decade, the number of metazoan, and particularly vertebrate, genomes has risen rapidly [12, 13]. However, generating the alignments of multiple genomes at the core of comparative genomics research is not trivial, and is made more difficult with increasing number, complexity and evolutionary distance between genomes [14]. Consequently, updates to whole-genome alignments (WGAs) have been relatively slow [15] and not reflective of the large number of available high-quality genome sequences. A group of species that are particularly under-represented in WGAs are marsupial mammals. Existing genomic alignments of 60, 100 and 144 vertebrate taxa frequently used in comparative genomics include the same three species and assemblies—gray short-tailed opossum (monDom5, released in 2007 [16]), tammar wallaby (macEug2, released in 2009 [17]) and Tasmanian devil (sarHar1, released in 2011 [18]). This reflects the historical sparsity of marsupial genomes, as until 2017 these were the only available. However, as of April 2025 there were reference genome assemblies for 27 marsupials available on NCBI. Yet these remain largely absent from most publicly available WGAs and can be challenging to incorporate into analyses.

Crucially, the range and number of taxa included in any comparative genomic study can profoundly affect inferences of constraint and divergence [19–23]. For example, initial genomic comparisons of the naked mole rat with 10 other mammalian species identified a single amino acid change in a protein associated with hairlessness (HR) [24]; this mutation was presumed to be phenotypically causal but subsequent investigation of the *HR* locus in 92 mammals found the same amino acid change in multiple other lineages that did not exhibit hairlessness [25]. Similarly, when studying the impact of phylogeny on inferences of convergence in marine mammals, Thomas et al. [26] found that as the number sampled taxa increased, the number of amino-acid substitutions called "convergent" decreased. A recent study by [27] evaluated human acceleration in 49 primate species and showed that almost two-thirds of HARs curated from previous analyses (which sampled the primate phylogeny more sparsely) were in fact conserved among primates.

Sparse taxonomic sampling also likely affected inferences of marsupial accelerated evolution in a previous study, Feigin et al. [28]. This work identified a set of conserved non-coding elements that were under sequence acceleration in both the marsupial thylacine (*Thylacinus cynocephalus*) and placental wolf (*Canis lupus*), known as thylacine and wolf accelerated regions (TWARs), and potentially underlying their well-characterised convergent craniofacial morphology [29, 30]. Limited by available resources, Feigin et al. [30] incorporated the thylacine genome to the 60-species WGA described above, which consists of 40 placental mammals, 3 marsupial mammals (plus the thylacine) and 17 other vertebrate taxa. In their analysis, Feigin et al. [28] identified almost 10 times the number of thylacine accelerated regions (10,910) than wolf accelerated regions (1,923).

Here, we re-assess the evidence for accelerated evolution in both the thylacine and wolf as well as multiple outgroup taxa in both clades, showing that the lack of marsupials genomes results in reduced power to accurately identify acceleration specific to each marsupial lineage. As a remedy, we have devised an approach that would allow inclusion of novel marsupial genomes into comparative analysis of vertebrate conserved elements. We present a workflow—Lift&Add—that takes advantage of two genome annotation lift-over tools, 1) UCSC liftOver, which relies on existing pairwise genome alignments between genome assemblies and 2) Liftoff, which maps genome annotations between assemblies with local alignment, as well as the sequence aligner MAFFT. We demonstrate and validate this workflow with a test dataset of alignments of conserved elements derived from the UCSC 60-way WGA of vertebrates, adding to this dataset 12 recently released marsupial genomes. We then assess the extent to which this additional phylogenetic breadth can enhance our study of lineage-specific acceleration in thylacine and other marsupials.

## Results

### Inferences of species-specific variation are limited by sparse taxonomic sampling

To investigate the impact of low phylogenetic coverage on estimations of thylacine and wolf accelerated evolution we repeated and extended the original analyses from Feigin et al. [28]. First, we re-characterised vertebrate conserved elements using an extension of the UCSC 60-way vertebrate WGA generated by Feigin et al. [28], to include the thylacine genome (the 61-way alignment, see Methods). Feigin et al. [28] limited their acceleration analyses to vertebrate conserved elements that intersected with craniofacial mouse ChIP-seq peaks from ENCODE; here, we use similar parameters but do not filter regions, assessing all vertebrate conserved elements for significant acceleration in the focal lineages (at a false discovery rate of 10%) (**Supplementary Table 1**). As in Feigin et al. [28], we find an excess of accelerated elements in the thylacine (16,698) relative to the wolf (11,861). Expanding our analysis to other taxa in the alignment, we consistently observe a higher number of accelerated conserved elements in marsupials than in placentals, with 18,278 conserved elements significantly accelerated in the Tasmanian devil (*Sarcophilius harrisi* and 21,763 in the tammar wallaby (*Monodelphis domestica*) but only 5,754 in the giant panda (*Ailuropoda melanoleuca*) and 11,202 in the domestic cat (*Felis cattus*).

However, overlap in accelerated regions was much higher among the marsupials—ranging between 24-44%—than among the placentals—ranging between 12-24% (Figure 1B). Of the 3.41 Mb of sequence that was accelerated in the thylacine, 44.0% was also accelerated in one or both of the other marsupials, which was significantly higher than the 12.2% of the wolf accelerated sequence shared with one or both of the other placentals (Binomial test *p*-value = 3.95 × 10*^−^*^323^, with expected shared proportion of 12.2%). This overlap could not be explained by evolutionary divergence. For instance, the tammar wallaby diverged 60-70 million years ago [31] from the other two marsupials and shared 24.1% of its accelerated sequence with them, an intersection size larger than that between each pair of placental species here. This overlap was also greater than that estimated between placental mammals with shorter divergence times in other studies [32, 33]. On the basis of these findings, we surmised that acceleration specific to individual marsupial taxa could not be reliably distinguished from marsupial-wide acceleration; further supporting this conclusion, only 22.4% of thylacine accelerated regions from the Feigin et al. [28] were reproduced in our analysis, compared to 88.3% of the wolf accelerated regions. This could be explained by the overall weaker signal for significant acceleration in the wolf relative to the thylacine (Supplementary Figure 1).

**Figure 1.**
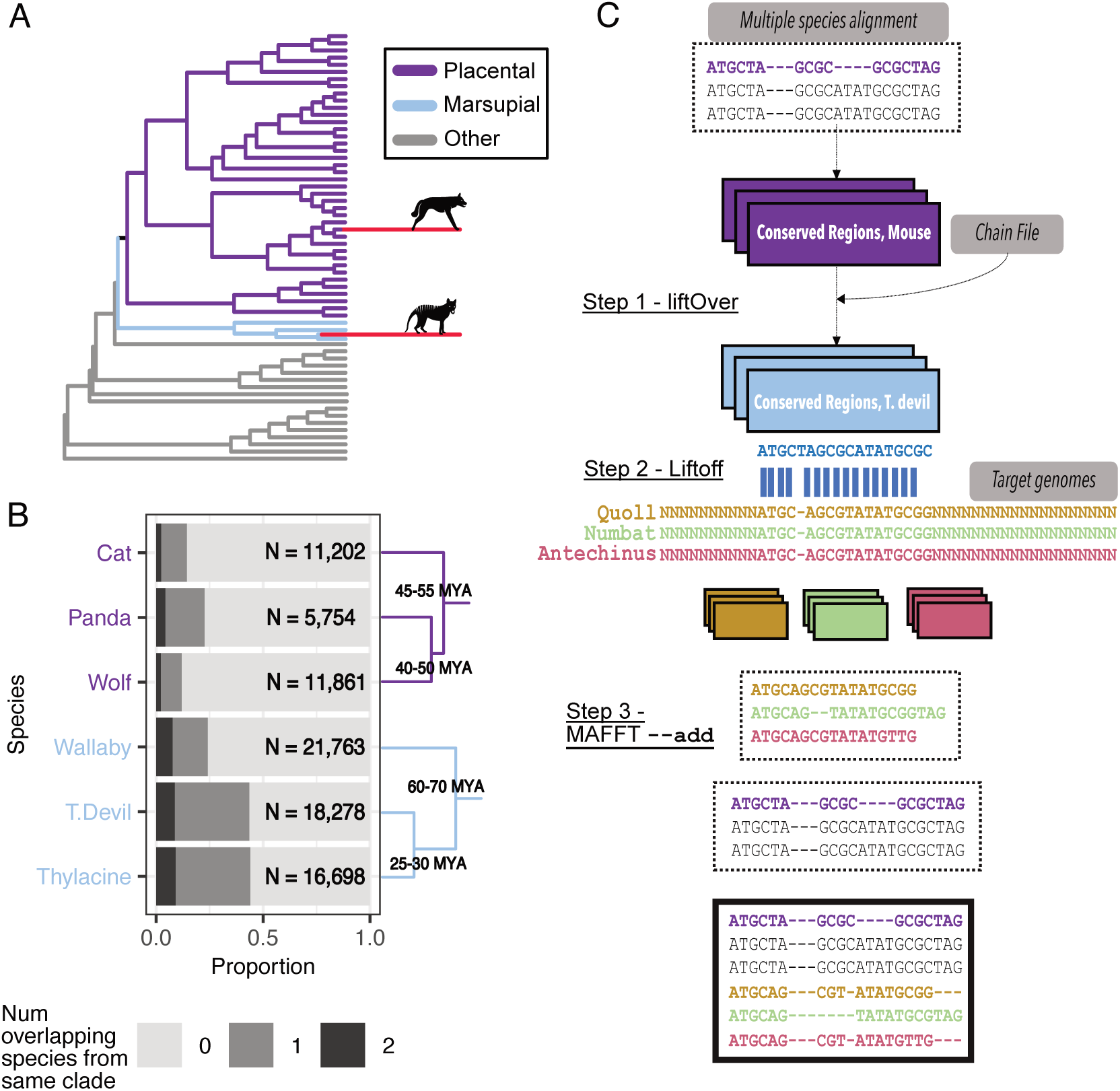
Increasing phylogenetic coverage for comparative genomic analyses. **A.** Phylogenetic overview of the 61 species used in Feigin et al. [28] and here to identify acceleration in the thylacine and wolf lineages (hypothetical acceleration visualised with extended branch lengths in red). **B.** Per-species proportion of total accelerated sequence intersecting with 0, 1, or 2 species from the same clade, i.e. with other marsupials or other placentals. **C.** Workflow to add sequences from new genomes to alignments of conserved elements. First, liftOver is used to map mouse coordinates of vertebrate conserved elements to marsupials with available chain files (here, the Tasmanian devil). Second, we map coordinates of vertebrate conserved elements between marsupials with Liftoff. The target genomes here are new marsupial assemblies that are not present in the whole-genome alignment or have chain files with mouse. Third, for each mapped element we add sequences from the newer marsupial genomes to the existing multiple-species alignments.

Reasoning that the dearth of marsupial taxa in the 61-way WGA was limiting our ability to identify thylacine-specific acceleration, we sought to develop a computational workflow that would allow us to incorporate newly available marsupial genomes into comparative analysis of vertebrate conserved elements. The workflow has been summarised in Figure 1C, and is described in detail in the following sections. Briefly, our approach, which we call Lift&Add, uses UCSC liftOver, Liftoff and MAFFT to allow for straightforward incorporation of genome sequences alignments at broadly conserved regions. Most existing whole genome alignments of vertebrates are referenced to a human or mouse genome assembly, as are conserved elements derived from these alignments. UCSC liftOver can move annotations between genome assemblies; however, it requires chain files, i.e. existing pairwise whole genome alignments between genomes. Liftoff on the other hand applies pairwise local alignment with minimap2 to map elements between a query genome and a target genome, an approach that works best between closely related taxa. Our workflow uses a combination of liftOver and Liftoff to map elements to new genomes in two steps, leveraging available chain files to minimise divergence times for Liftoff [34]. Mapped elements are subsequently compiled and added to conserved sequence alignments with MAFFT [35]. The final output consists of new alignments for vertebrate conserved elements covering an expanded breadth of taxa.

### Optimising Liftoff parameters to map short elements over long evolutionary distances

Existing documentation and published use of Liftoff is limited to genes, exons and transcripts. Thus we first sought to evaluate Liftoff’s performance with our mostly non-coding dataset by comparing its performance to that of to UCSC liftOver, which requires the use of a chain file to map coordinates between genome assemblies or species. In order to create a test dataset to evaluate Liftoff with, we began by identifying 17,155 discrete vertebrate conserved elements >50 bp aligning to mouse chromosome 19 in the original 60-way WGA (without the thylacine) with PHAST. Annotating our dataset against mm10, which the WGA is referenced to, we determined that 28.0% of elements (4,796) overlapped protein coding sequences (CDS), 5.5% (949) 3’ or 5’ untranslated regions (UTRs) and the remaining 66.5% (11,410) were intergenic or unannotated. We then used liftOver and existing chain files to map 13,178 of these elements from mouse (mm10) to the opossum (monDom5) (Supplementary Figure 2, **Supplementary Table 2**).

To evaluate the performance of Liftoff on this heterogenous test dataset, we then mapped these elements from the opposum (monDom5) to the Tasmanian devil (sarHar1) either liftOver or Liftoff, and compared the results (Figure 2A; [31]). Given the long divergence time between these taxa (TMRCA ≈80-90 million years), substantially fewer elements mapped with Liftoff (5,699 of 13,178, 43.2%) than with liftOver (11,207, 85.0%) (Figure 2B). Of the 5,469 elements successfully mapped with both liftOver and Liftoff, 5,448 had overlapping output coordinates between the two methods (with an average 99.0% of the liftOver output bases overlapping with Liftoff output bases). Similar proportions of protein-coding (44.2%), UTR (42.4%) and intergenic/unannotated (51.4%) elements were successfully mapped with Liftoff (**Supplementary Table 3**); however Liftoff performed significantly worse with shorter elements (mean size (bp) mapped elements 200.5, unmapped elements = 104.3; pairwise t-test *p*-value *<* 2.2 × 10*^−^*^16^) (Figure 2C).

**Figure 2.**
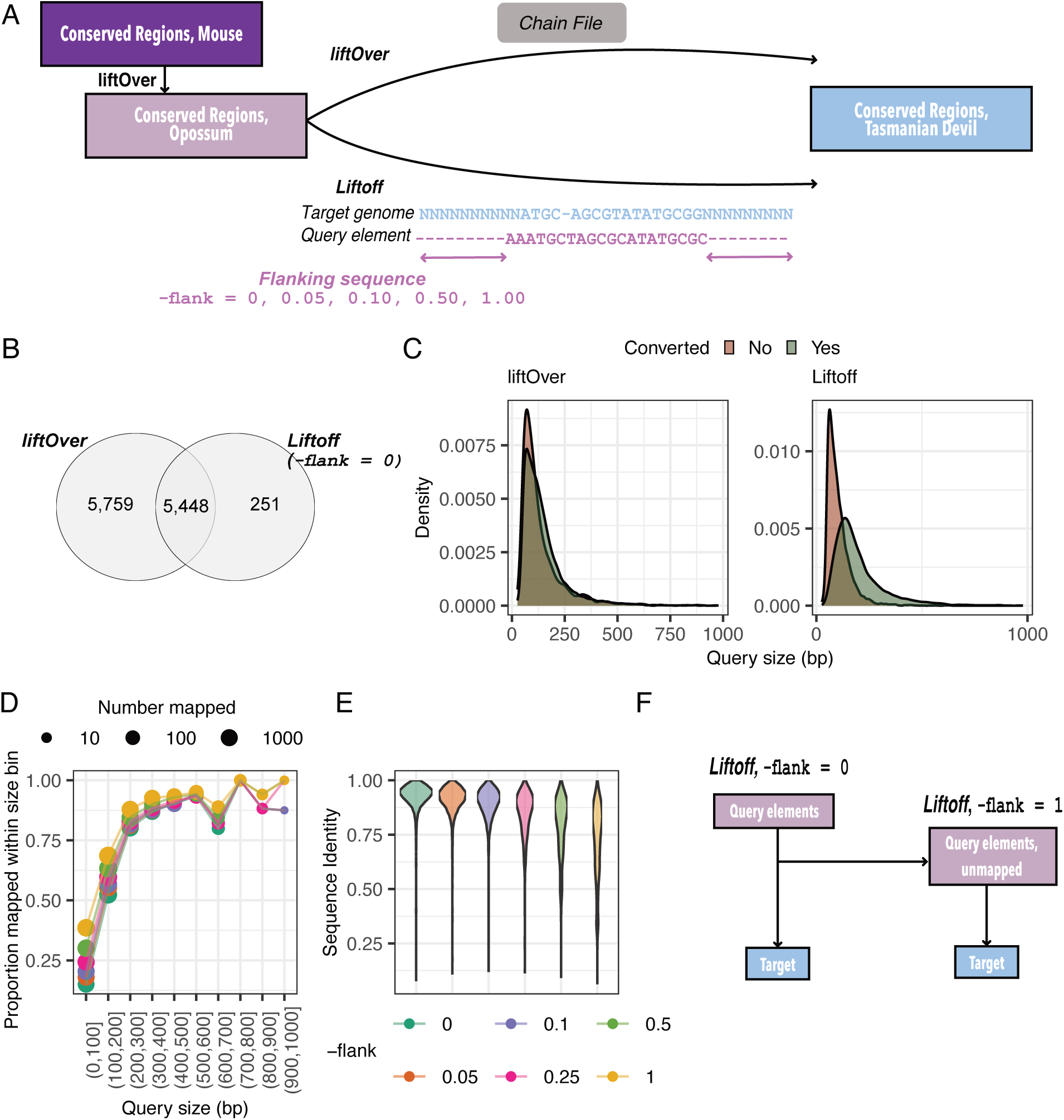
Testing Liftoff. **A.** Coordinates for vertebrate conserved elements were mapped from opossum to Tasmanian devil with both liftOver and Liftoff. **B.** Overlap between output from liftOver and Liftoff. **C.** Distribution of sizes of elements that did and did not successfully map from opossum to devil with liftOver and Liftoff. Elements greater than *>* 1000 bp in length are excluded. **D.** Proportion of opossum query elements in each 100 bp size interval successfully mapped at different thresholds of -flank. **E.** Sequence identity between query opossum and target Tasmanian devil elements at different threshold of -flank. **F.** Final Liftoff strategy used in our workflow.

Liftoff allows flanking genomic sequence to be symmetrically added to query elements (parameter -flank, default 0). We found that increasing the total amount of flanking sequence added stepwise (-flank = 0.05, 0.10, 0.25, 0.5 and 1, where these values are multiples of the original query length) consistently increases the number of elements mapped (Supplementary Figure 3A, **Supplementary Table 4**). Elements only mapping at -flank *>* 0 were significantly shorter than those mapped at -flank = 0 (pairwise t-test *p*-value *<* 2.2 × 10*^−^*^16^), demonstrating that adding flanking sequence improves Liftoff performance for shorter elements, primarily for those between 0-200 bp (Figure 2D, Supplementary Figure 3B). However, genomic sequence flanking vertebrate conserved elements is likely to be under less constraint itself and therefore may reduced alignment specificity. Indeed, with added flanking sequence the average alignment coverage and sequence identity between the aligned query and target elements decreases, and the number of elements that are partially mapping (i.e. with alignment coverage and/or sequence identity =*<* 0.5, as defined by Liftoff) increasing from 44 at -flank = 0 to 806 at -flank = 1. (Figure 2E, Supplementary Figure 3C, Table **Supplementary Table 4**)

To minimize partial mapping, we tested a successive approach, whereby we first use Liftoff to map elements between query and target with no flanking sequence (-flank = 0) (Supplementary Figure 4). Subsequently, for sequences that do not map/partially map, we repeat Liftoff ("re-map") with -flank = 1 to maximize recovery. Initial Liftoff maps 5,699 elements (43.2% of the opossum dataset) to Tasmanian devil genome. An additional 2,344 elements (31.3% of previously unmapped elements) re-map with the inclusion of flanking sequence. Only 191 elements displayed partial mapping (0.02% of 7,999); excluding these, the total number of successfully mapped elements was 7,805 (59.2% of the query dataset). This was higher than a single round of Liftoff at any threshold of -flank (**Supplementary Table 4**). Of these 7,805 elements, 7,370 have overlapping coordinates with the output of liftOver from opossum to the Tasmanian devil (average overlap 99.2%), 391 were mapped with Liftoff but not liftOver and 44 display disagreement between the two methods.

### Mapping vertebrate conserved elements to 12 new marsupial genome assemblies

Our results with the Tasmanian devil and opossum suggested that Liftoff could be used to map these vertebrate conserved elements (both coding and non-coding elements) between more closely-related marsupial taxa, with the ultimate goal—in this case—of re-evaluating thylacine accelerated evolution. Focusing still on conserved elements extracted from mouse chromosome 19, we used UCSC liftOver to map the conserved elements to the two marsupial genomes with publicly available mm10 chain files—the Tasmanian devil (sarHar1/Devil_ref_v7.0) and tammar wallaby (macEug2/Meug_2.0) Supplementary Figure 2, **Supplementary Table 2**). We then curated 10 new genome assemblies of species from 8 marsupial families not represented in the UCSC 60-way WGA of vertebrates (including the newer thylacine genome assembly [36]) as well as newer assemblies for Tasmanian devil (mSarHar1.11, 2022) and tammar wallaby (mMacEug1, 2021) (**Supplementary Table 5**) [37]. The older tammar wallaby assembly (macEug2 [17] was used as the Liftoff query genome for the six new marsupials within Diprotodontia, while the Tasmanian devil (sarHar1) [18] was used as query genome for the five marsupials within Dasyuromorphia. Our final taxa, the monito del monte (*Dromiciops gliroides*, mDroGli1) is the only extant species in the order Microbiotheria, and equally divergent from Tasmanian devil and tammar wallaby (TMRCA 60-70 million years) [31]. We used the Tasmanian devil genome (sarHar1) as query genome in this instance due to its higher contiguity and completeness relative to the tammar genome (macEug2) [17, 18, 38].

Starting with the same set of mm10 chromosome 19 elements as above, we mapped with liftOver 12,345 and 10,692 to the Tasmanian devil (sarHar1) and tammar wallaby (macEug2) genomes, respectively (Table 1). With Liftoff, we then mapped all 12,345 vertebrate conserved elements from Tasmanian devil to one or more of its target genomes, and 10,688 (99.9%) conserved elements from tammar wallaby to one or more of its target genomes (Figure 3A and B, Supplementary Figure 5A-D). For both groups, a majority of the elements mapped to all six species—8,564 (69.4%) from the Tasmanian devil and 8,098 (75.7%) from the tammar wallaby. Elements that mapped to all new target species from the Tasmanian devil or the tammar wallaby were significantly longer than those did not, consistent with size being a primary driver of success with Liftoff (pairwise t-test Dasyuromorphia: *p*-value= 1.386 × 10*^−^*^5^, Diprotodontia: *p*-value= 8.718 × 10*^−^*^145^ (Supplementary Figure 5E and F). No bias was observed in the type of elements—CDS, UTR or intergenic/unannotated—that were successfully mapped (Figure 3C and D).

**Figure 3.**
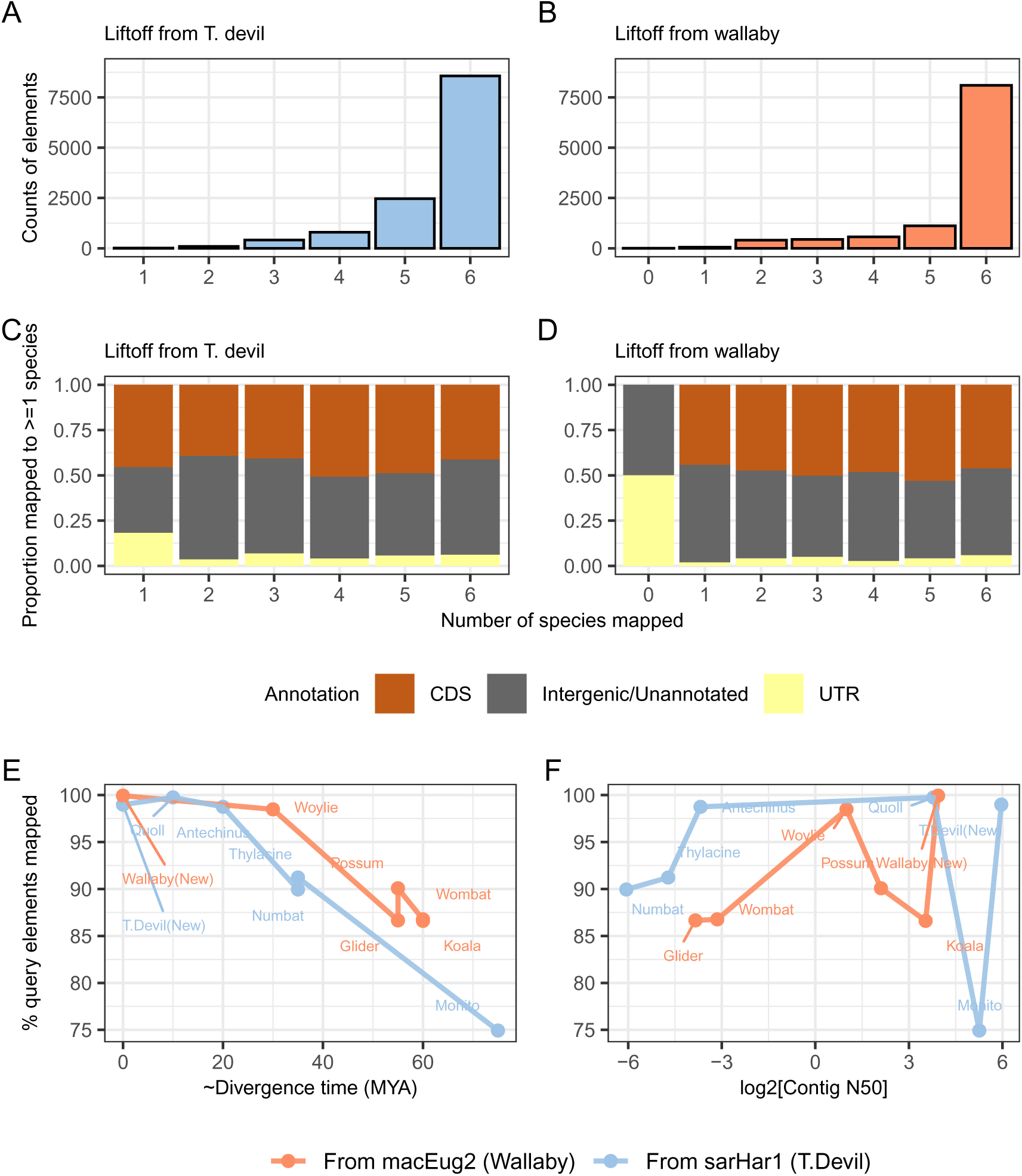
Mapping conserved elements from Tasmanian devil (sarHar1) and tammar wallaby (macEug2) to target marsupial genomes. **A. and B.** Counts of query elements that were mapped to 0 or more species. **C. and D.** Annotations for query elements that mapped to 0 or more species. **E.** Estimated divergence time versus percentage of query elements mapped. **F.** Genome contiguity (Contig N50) versus percentage of query elements mapped.

**Table 1.**
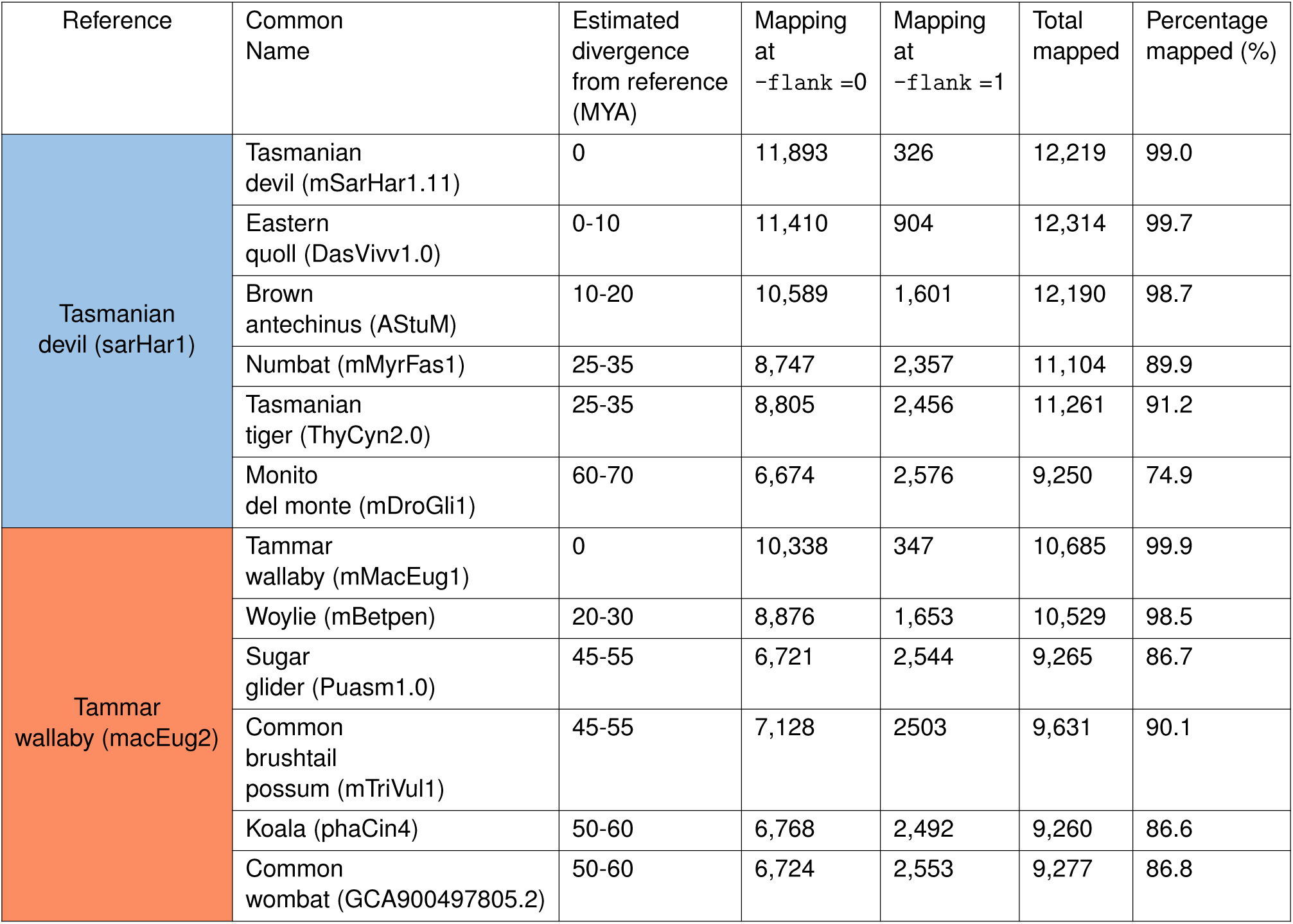
Liftoff results for coordinate conversion from the Tasmanian devil/tammar wallaby to 12 new marsupial assemblies.

The percentage of elements mapped was highly negatively correlated with the evolutionary distance between query and target genome (Spearman’s *ρ* -0.95 (*p*-value = 1.46 × 10*^−^*^6^) (Figure 3E). Nearly all query elements mapped across different assemblies of the same species (98.9% from sarHar1 to the mSarHar1.11 for Tasmanian devil, and 99.9% from macEug2 to mMacEug1 for tammar wallaby), whereas the lowest percentage of mapped elements was between the distantly related Tasmanian devil and monito del monte (74.9%; estimated divergence time ∼ 60-70 MYA). Additionally, the proportion of elements that could be mapped with -flank = 1 was correlated positively with divergence time (Spearman’s *ρ* = 0.98, *p*-value = 3.181 × 10*^−^*^8^), confirming that as sequence divergence between query and target increases, adding flanking sequence and increasing query length has a greater impact on local alignment quality (Supplementary Figure 5G). Lastly, there was no overall association between contig N50 and percentage of query elements mapped (Spearman’s *ρ* = -0.19, *p*-value = 0.54), likely due to the short sizes of the query vertebrate conserved elements (query element mean length, Tasmanian devil (sarHar1): 161.9bp, tammar wallaby (macEug2) 147.2 bp) (Figure 3F).

Aggregating results from the two clades, we successfully mapped 13,297 vertebrate conserved elements from Tasmanian devil (sarHar1) and tammar wallaby (macEug2) to at least one of 12 new genomes. Excluding the newer assemblies for Tasmanian devil (mSarHar1.11) and tammar wallaby (mMacEug1), 13,288 (99.9%) mapped to at least one of 10 new genomes (Supplementary Figure 6). Of these, 5,986 (45.0%) mapped to all 10 species. Only 77% of the 17,155 of vertebrate conserved elements from mouse chromosome 19 mapped to at least one new marsupial genome (excluding newer assemblies); considering only alignment blocks that contain a marsupial (14,425), we were able to map the vast majority (92.1%) to new marsupial genomes.

### Lift&Add results correlate with inter-species karyotypic relationships

As a means of confirming the accuracy of our approach, we examined the genomic order of mapped elements in the new target genomes. Karyotypes are highly conserved across species in Dasyuromorphia (chromosome number 2n=14) [39]. Thus, we expected the genomic order of mapped elements to be co-linear between the query Tasmanian devil genome and the target Dasyuromorphia genomes. However, the query Tasmanian devil (sarHar1) genome assembly was only scaffold-level. Instead, we assessed co-linearity between the newer chromosome-level Tasmanian devil genome assembly (mSarHar1.11) to the other Dasyuromorphia target genomes, noting that since our test dataset includes only conserved elements in mouse chromosome 19, we do not expect to cover the whole genome in any species. As expected, we observed near-perfect co-linearity in genomic order of conserved elements (Spearman’s *ρ* = 0.98, *p*-value *<* 2.2 × *e^−^*^16^) between the closely-related Tasmanian devil and quoll genomes (divergence time ≈5-10 MYA) [39] (Figure 4A). A perfect inversion in the order of elements from chromosome 6 when comparing between mSarHar1 and the Eastern quoll (DasVivv1.0) could be indicative of structural variation but could also originate from differences in chromosome assembly and annotation.

**Figure 4.**
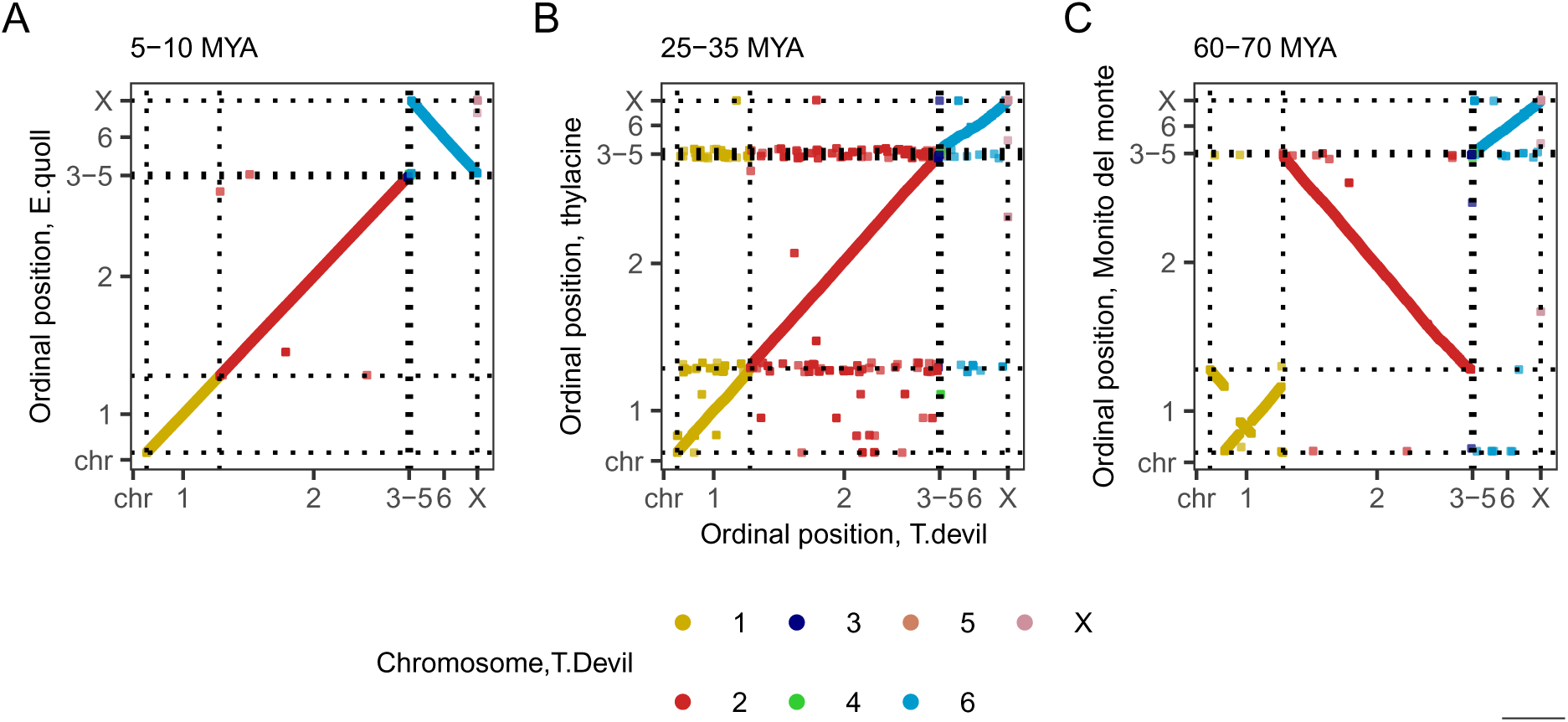
Correlation in genomic order of mapped elements between. **A.** Tasmanian devil (mSarHar1.11) and Eastern quoll (DasVivv1.0), **B.** Tasmanian devil and Thylacine (ThyCyn2.0) **C.** Tasmanian devil and Monito del monte (mDroGli1). Elements are coloured by their chromosomal location in the Tasmanian devil

Comparing the results of mapping to the new Tasmanian devil assembly (mSarHar1.11) and the more distantly related extinct thylacine (divergence time ∼ 25-35 MYA) we again found a high degree of co-linearity of mapped elements (Spearman’s *ρ* = 0.96, *p*-value *<* 2.2 × *e^−^*^16^) (Figure 4). However, there are more elements that show discordant genomic order between the new Tasmanian devil genome and the thylacine genome. The thylacine chromosome-level assembly (ThyCyn2.0, 2022) was derived from short contigs (contig N50 0.015 Mb) that were assembled into larger scaffolds with reference to mSarHar1.11 [36]; as such we speculate that this discordance was likely due to instances of mis-assembly of the thylacine genome due to the challenging nature of the original data. By examining the coordinates of these conserved elements between the two genomes (Supplementary Figure 7), we find support for this theory, with three clusters of elements from likely repetitive regions in the Tasmanian devil mapping to variable positions in the thylacine.

Lastly, Spearman’s correlation in the genomic order of mapped elements between the Tasmanian devil and distantly related monito del monte (which also has a karyotype 2n=14) was 0.52 (*p*-value *<* 2.2× *e^−^*^16^) (Figure 4C). Elements from chromosome 1 and chromosome 6 display linear agreement in order; a near-perfect inversion of coordinates for elements in chromosome 2 lowered the overall correlation. As with the quoll comparison, this likely reflects differences in genome assembly.

As with the Tasmanian devil, the older tammar wallaby assembly (macEug2) was also scaffold-level, but the *newer* assembly (mMacEug1) was chromosome level; we thus again compared order of mapped elements in this new assembly to their order in the brushtail possum genome (mTriVul1), the only other Diprotodontia chromosome-scale assembly. The Diprotodontia taxa display high variability in karyotypes with several instances of chromosomal reshuffling in their evolutionary history, particularly in the Macropodiformes, and the brushtail possum has two more chromosomes (2n=20) than the tammar wallaby (2n=16) (TMRCA 45-55 MYA) [39, 41]). Overall, we observed a strong (but negative) correlation in genomic order between species (Spearman’s *ρ* = -0.93, *p*-value *<* 2.2 × *e^−^*^16^) (Figure 5A), driven primarily by strong collinearity between tammar wallaby chromosomes 1 and 2 and brushtail possum chromosomes 8 and 6 respectively, although tammar chromosome 1 and possum chromosome 8 are inverted relative to each other. However, results for elements mapped to tammar wallaby chromosomes 3 or X were more variable. To better understand their complex cytogenetic relationship, we referred to chromosome painting results from [40] (Figure 5B, Supplementary Figure 8, **Supplementary Table 6**). In this work, [40] showed that chromosome 2 of the tammar wallaby hybridized to chromosomes 6 and 3 of the brushtail possum, aligning with our observation that 95.9% of elements from tammar wallaby chromosome 2 mapped to chromosome 6 of the brushtail. Chromosome 3 of the tammar wallaby hybridized to brushtail possum chromosomes 1, 5 and 9 in the original chromosome painting results; in our data 91.4% of tammar wallaby chromosome 3 elements mapped to brushtail possum chromosome 1 and an additional 6.2% mapped to brushtail possum chromosome 5. However, chromosome painting results results suggested that tammar wallaby chromosome 1 is alignable to chromosome 4, 7 and 9 of the brushtail possum, in our data the majority (98.4%) of elements from tammar wallaby chromosome 1 map to either chromosome 8 and 9 of the brushtail possum. Looking closer, we found that chromosome 7 and 8 of the brushtail are 275 and 267 Mb respectively, and likely difficult to differentiate cytogenetically. Altogether, we were able to show that our results agreed with expectations of both karyotype conservation in Dasyuromorphia and karyotype rearrangement in Diprotodontia.

**Figure 5.**
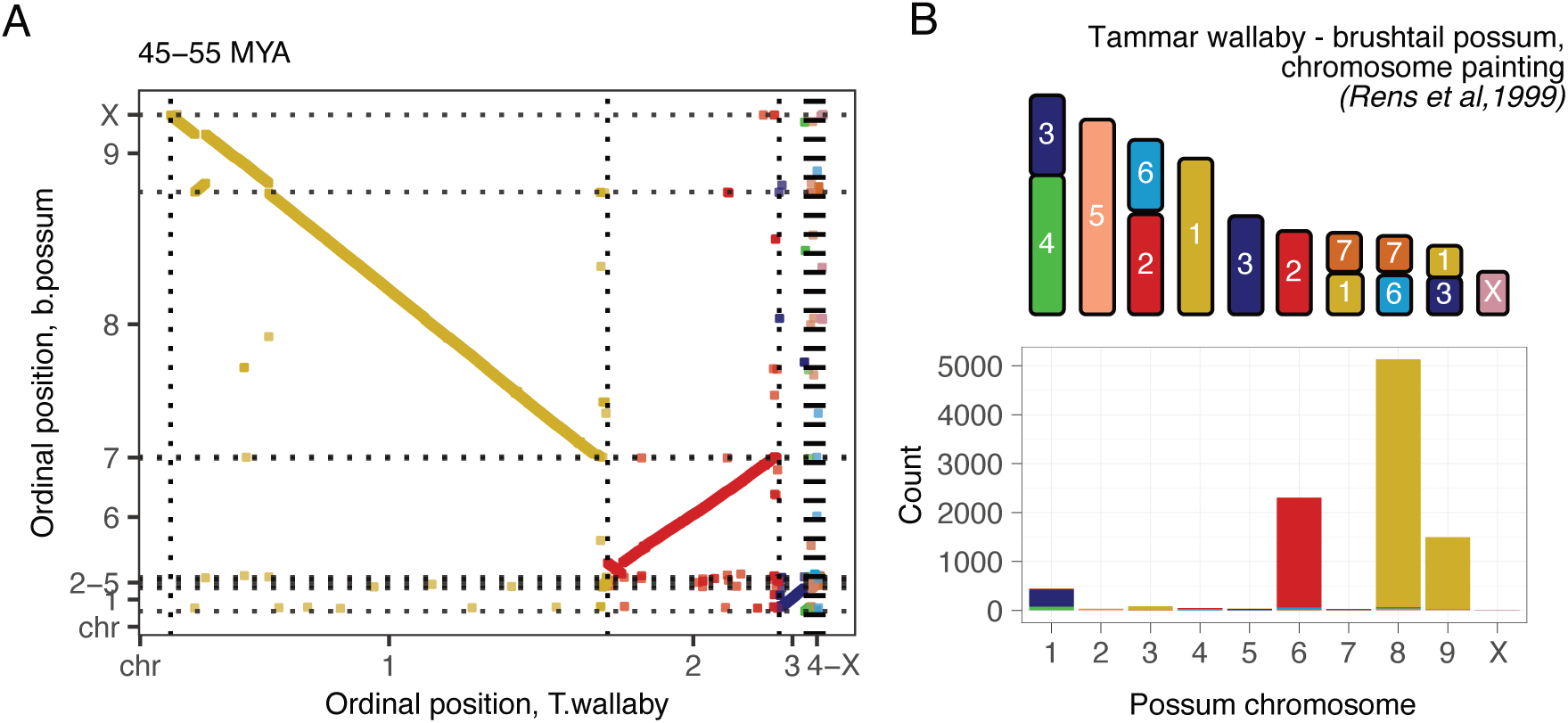
Tammar wallaby and brushtail possum. **A.** Correlation in genomic order of mapped elements between tammar wallaby (mMacEug1) and brushtail possum (mTriVul1). Elements are coloured by their chromosomal location in the tammar wallaby. **B.** Relationship between the tammar wallaby and brushtail possum karyotypes, as determined by [40], and counts of test elements mapping to each brushtail possum chromosome, coloured by their location in the tammar wallaby genome.

### Generating new multiple-species alignments with MAFFT

Having mapped vertebrate conserved elements to new marsupial genomes, the next step in our pipeline is to incorporate the relevant sequences back to the corresponding multiple-species alignments with MAFFT –add, as a prerequisite for testing for acceleration. An additional parameter with MAFFT, –keep-length allows the structure of the original alignment to be preserved when new sequences are added: if the new marsupial sequences are longer than the alignment, MAFFT trims the unaligned bases, such that only the original sequence is retained, controlling for the inclusion of flanking sequence in the steps above.

To evaluate the resulting alignments, we leveraged sequences from existing marsupials in the 60-way WGA, reasoning that they should align well with added marsupial sequences (Figure 6). For every alignment of a conserved element, we calculated the pairwise alignment distance (proportion of sites that differ between two sequences in an alignment) between the Tasmanian devil (sarHar1) or tammar wallaby (macEug2) sequences and each new marsupial sequence (Figure 6B and C). Overall, the sequences of the added Dasyuromophia species were significantly more similar to Tasmanian devil (sarHar1) than sequences from the Diprotodontia species or the opossum (**Supplementary Table 7**). Conversely, sequences of all added Diprotodontia species were significantly more similar to tammar wallaby (macEug2) than sequences from the Dasyuromorphia species, the monito del monte or the opossum (**Supplementary Table 8**). For each species, the mean pairwise distance from the Tasmanian devil was positively correlated with divergence time (Spearman’s *ρ* 0.80 (*p*-value 0.0009), as was the mean pairwise distance from the tammar wallaby (Spearman’s *ρ* 0.86 (*p*-value = 7.31 × 10*^−^*^5^) (Figure 6D and E). These observations suggested that the new alignments were largely congruous with phylogenetic expectations.

**Figure 6.**
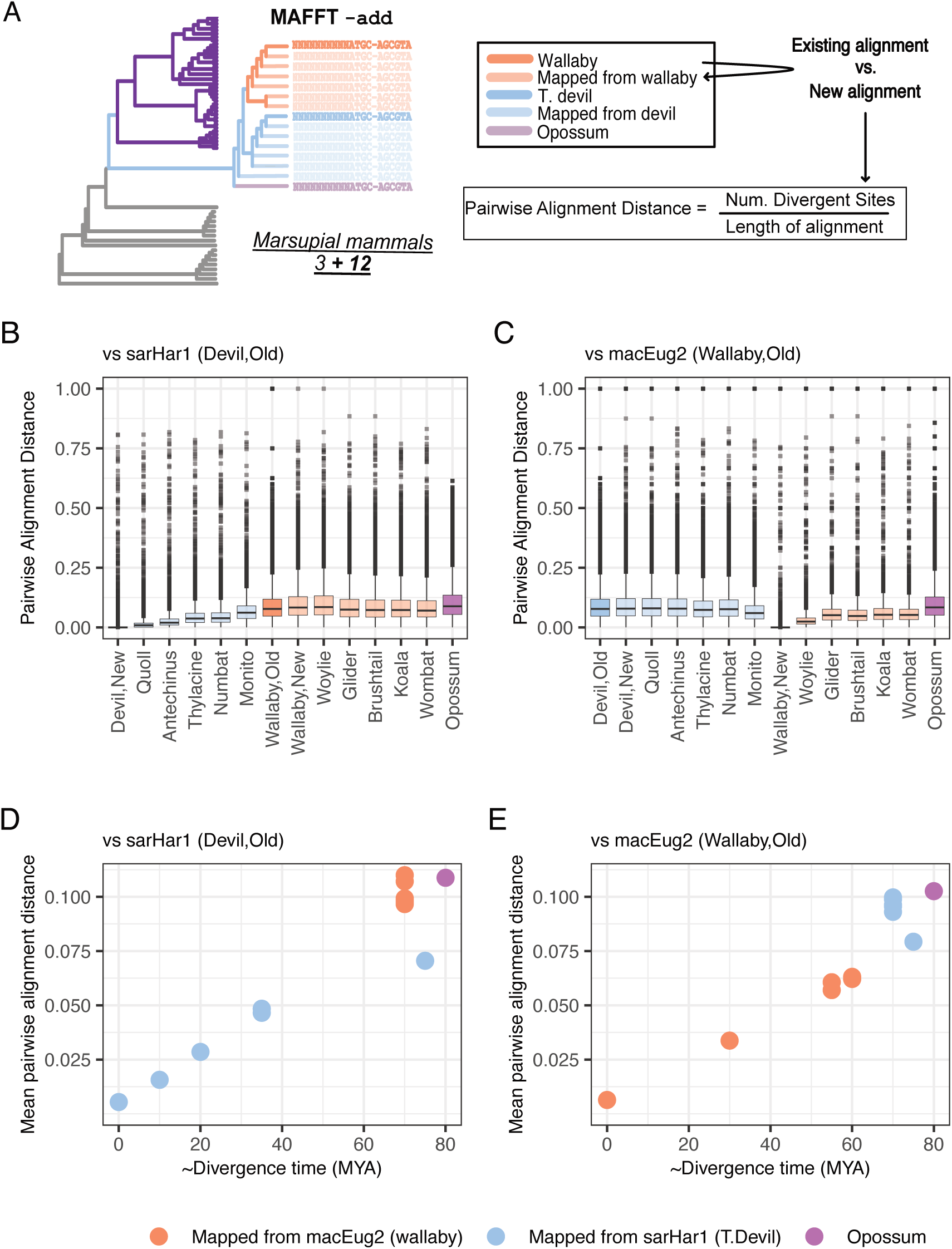
Alignment with MAFFT. **A.** MAFFT –add was used to incorporate sequences from 12 new marsupials genomes to existing sequence alignments for vertebrate conserved elements. To QC the resulting alignments, we calculated the pairwise alignment distance between existing and added sequences. Pairwise alignment distance between **B.** the Tasmanian devil and each new and old marsupial genome and **C.** the tammar wallaby and each new and old marsupial genome. **D.** Estimated divergence time from the Tasmanian devil versus average pairwise distance (across all elements) from the Tasmanian devil. **E.** Estimated divergence time from the tammar wallaby and versus average pairwise distance from the tammar wallaby.

### Comparing relative rates of evolution with RERConverge

Lastly, we examined how output alignments from our Lift&Add workflow can be explored for variations in evolutionary rates, using here the R package RERConverge to revisit thylacine and wolf acceleration [42]. We added the gray wolf (mCanLor1.2 released 2021, contig N50 34.4 Mb and scaffold N50 65.8 Mb [43]) to our alignments with Lift&Add, mapping the test dataset of 17,155 conserved elements from the mm10 chromosome 19 to the dog genome (canFam3) with UCSC liftOver, followed by Liftoff of elements from the dog genome to the target gray wolf genome. Of the total 17,067 (99.9%) vertebrate conserved elements that mapped to the wolf genome, 13,200 also mapped to at least one marsupial genome, excluding 9 elements only mapping to the new Tasmanian devil assembly (mSarHar1.11) and/or tammar wallaby assembly (mMacEug1). With MAFFT, we added conserved sequences from the wolf to the multiple sequence alignments of each of these 13,200 elements. Our final alignments contained 13 new genomes: this new chromosome-level assembly of the gray wolf and the 12 marsupial genomes detailed above.

RERConverge is specifically designed to correlate convergent phenotypic traits (binary or continuous data) with shifts in evolutionary rates [42]. As in the original Feigin et al. [28] study, we here looked at thylacine-wolf convergence as a binary trait (coded as 1 for thylacine and wolf, and 0 for all other taxa). We first estimated individual species trees for each conserved element, which were then averaged to derive a single consensus tree, describing the overall evolutionary rate across all elements for each branch [42] (Supplementary Figure 9). Comparing this consensus tree to the phylogenetic model for neutral evolution derived from the original 60-way WGA [37], we observed positive correlation in phylogenetic distances for each pair of taxa (Spearman’s *ρ* 0.98 (*p*-value *<* 2.22 × *e^−^*^16^). As the RERConverge tree was derived from conserved genomic regions, these pairwise distances are on average 0.22 times shorter those in the neutral tree.

After estimating the consensus tree, we then calculated element-specific rates of evolution—the relative evolutionary rates (RERs), where a positive value is indicative of acceleration and a negative one of constraint—for every element in our test dataset [42, 44]. This included elements overlapping (with between 47% to 100% overlap) six thylacine and wolf accelerated regions (TWARs) from Feigin et al. [28] which aligned to mouse chromosome 19 (Figures 7A-F). By using Lift&Add we were able to incorporate between 4 and 9 additional marsupial genomes to these TWAR elements. Both the thylacine and wolf had positive RERs for all six TWAR-corresponding elements. This confirms accelerated evolution of these elements in the thylacine and wolf, as was previously determined in Feigin et al. [28].

**Figure 7.**
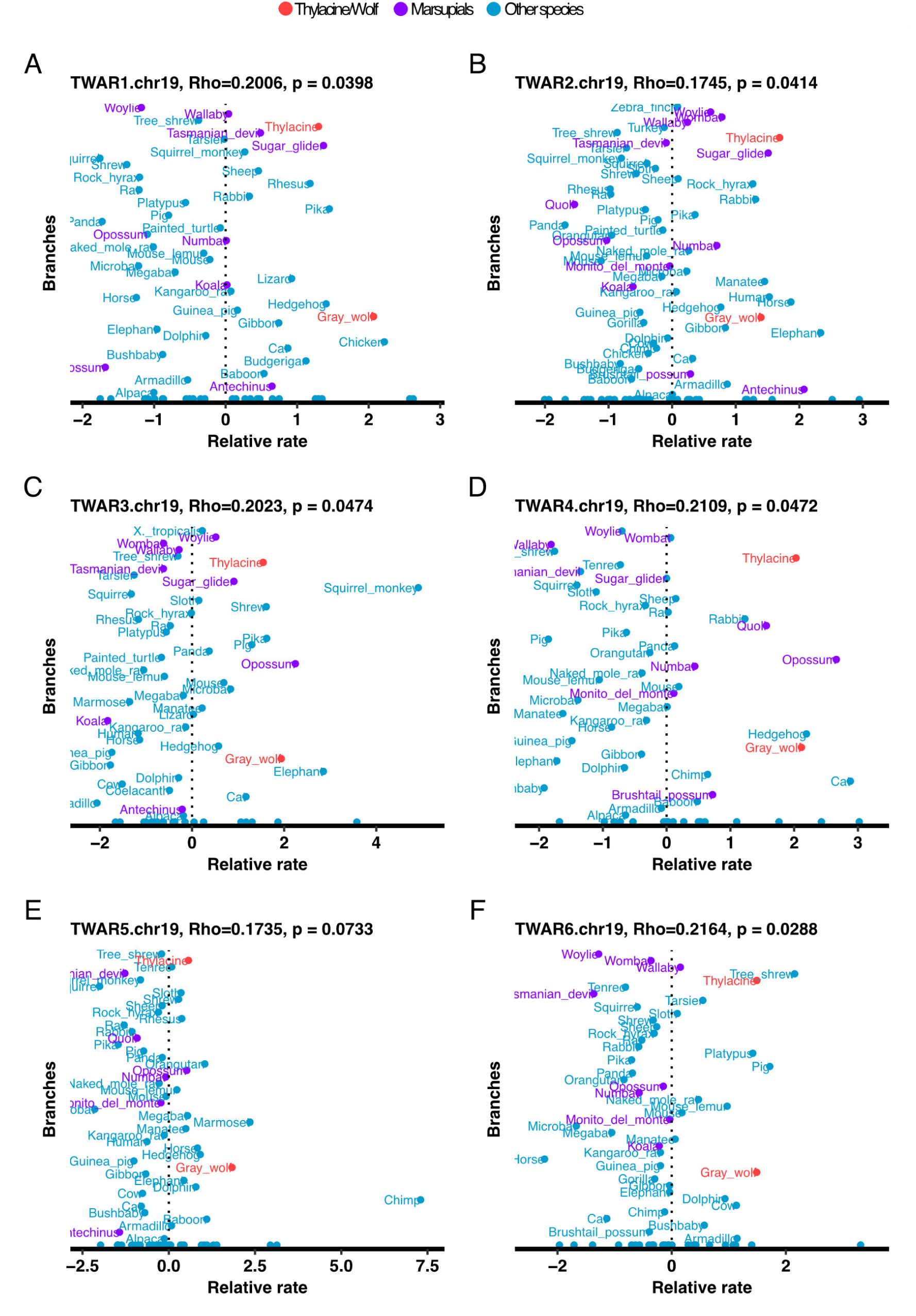
Relative evolutionary rates and phenotype association output (Rho and *p*-values) for elements overlapping TWARs. Rho is the correlation between the relative evolutionary rate and the convergent phenotype and P is the nominal *p*-value for this association.

However, multiple other marsupial taxa are also under accelerated evolution for these TWAR-corresponding elements. In elements equivalent to TWAR 1, 2, 3 and 4 from chromosome 19, the thylacine is not the marsupial species with highest RER (Figures 7A-D, marsupials highlighted in purple). In fact, multiple other marsupials have positive RERs greater than or comparable to the thylacine. These results highlight the importance of densely sampling more closely related taxa in comparative genomic analyses for specific identification of lineage-specific variation. Across 8,505 conserved elements aligned to at least 20 vertebrate species, we were unable to identify any elements where RERs were correlated with the thylacine-wolf binary convergence (at a FDR of 5%), including any of the six TWARs included in this re-analysis, (Supplementary Figure 10).

## Discussion

The power of comparative genomics analyses to resolve evolutionary constraint and acceleration is a function of the sequence diversity that is sampled [19, 20, 23, 25]. Here, we show that the paucity of marsupial genomes used in Feigin et al. [28] and our own re-analysis limited investigation of thylacine-specific acceleration in vertebrate conserved elements. Subsequently, we presented a bioinformatic workflow, Lift&Add, that 1) robustly maps vertebrate conserved elements to new genome assemblies and 2) can add sequences of conserved elements from these new genomes back to existing alignments, enabling more rigorous inter-sequence comparisons.

The 61-way WGA used in the [28] as well as in our analysis here has 41 placental mammals (representing 40 of the 127 of the extant placental families), 4 marsupial mammals (representing 4 of the 22 extant marsupial families) and 17 other vertebrates. Acceleration is defined relative to other taxa [45, 46]; given the biased distribution of species, any variation that is common to all marsupials will nonetheless be depleted among the compared species. Furthermore, while broad comparisons over long evolutionary distances have been historically instrumental in identifying highly conserved genomic segments (e.g., human and mouse), comparisons with closely-related taxa (e.g., human and chimpanzee) provide more accurate inferences of lineage-specific sequence evolution. Due to these taxonomic limitations, we observed much higher estimates of accelerated regions in the three marsupials—thylacine, Tasmanian devil, and tammar wallaby—than in the three placentals—wolf, giant panda, and the domestic cat. At the same time, we also observed a much greater overlap in accelerated regions among the marsupials than the placentals, pointing to a lack of lineage specificity.

Defining lineage-specific acceleration accurately in these species necessitates inclusion of more marsupials. Generating new WGAs is computationally expensive, as is adding multiple genomes to an existing alignment. Instead, we hypothesized that Liftoff [34] could be feasibly used to map both exonic and non-coding vertebrate conserved elements between genome assemblies, considering that these vertebrate conserved elements 1) are defined by alignment and a high degree of sequence similarity across distant taxa, 2) are on average shorter than most exons 3) and thus require fewer structural considerations for alignment than genes and transcripts. Success with local alignment is greatest over shorter evolutionary distances; we thus adopted a two-step approach, using an initial liftOver step to map the elements in mouse coordinates to query marsupial genomes. As expected, the percentage of mapped elements with Liftoff was negatively correlated with divergence distance from the query genome. Yet, even at the longest evolutionary distance between query and target genome (60-70 MYA, between the Tasmanian devil and monito del monte) 75% of conserved elements remained mappable. Crucially, we observed no bias for successful mapping of protein-coding or non-coding elements. Examining positional information of mapped elements between genomes (as in the example given by Shumate and Salzberg [34]), we observed concordance with expected cytogenetic relationships between taxa.

The biggest predictor of successful mapping with Liftoff was the length of query elements. Addition of flanking sequence to short (50-200 bp) conserved elements was associated with improved mapping. The sequence aligner used by Liftoff, minimap2, can map short reads but displays better performance with longer reads and has a minimum recommended read length of 100 bp [47]. This could underlie the poor performance of Liftoff for shorter elements. It can be argued that adding flanking sequence to a vertebrate conserved element introduces non-conserved bases, promoting spurious mapping. However, by comparing Liftoff and liftOver output, we confirmed that our two-step approach displays comparable accuracy with liftOver. Furthermore, the ability to align a conserved element across species decays proportional to the phylogenetic distance between them [3, 48]. As such, even if genomic sequence flanking a query element was not determined to be conserved across vertebrates, it is likely to be more conserved among just the marsupial species. We exclude elements that map with sequence identity lower than 50% of the query element, as recommended by Liftoff Shumate and Salzberg [34]. While reducing incorrect mapping, this limits investigation of sequence variation unique to the added taxa. Elements that are excessively divergent in the query or target genomes, such as those containing large deletions or insertions, will not be mapped. As such, our Lift&Add workflow is best suited to add phylogenetic context to a multiple species alignment or to investigate more subtle sequence changes in the added taxa (which will also likely be more frequent in highly constrained genomic regions).

Finally, we present here an example of an investigation that can be carried out following the use of Lift&Add, using RERConverge to explore evolutionary rates in our test dataset to re-evaluate the evidence for convergent acceleration between the gray wolf and the thylacine. No element in this dataset had evolutionary rates significantly associated with the thylacine-wolf convergent phenotype. We reanalysed six TWARs from Feigin et al. [28] within the context of additional marsupial sequences. Encouragingly, all six were accelerated both in thylacine and the wolf. Four of these, however, also had positive relative rates in other marsupial species. Although narrow, this analysis indicated the importance of dense sampling from closely-related taxa in the study of lineage-specific acceleration. Interestingly, for each of the six TWAR-corresponding elements, diverse subsets of placental mammal also displayed comparable acceleration to the wolf. This potentially also highlights the challenge of encoding the complex craniofacial phenotype observed in the thylacine and wolf as a binary trait.

## Materials and Methods

### Re-analysis of thylacine and wolf accelerated elements

For our initial re-evaluation of thylacine-wolf acceleration, we obtained the 61-way WGA, referenced to the mouse mm10 genome, from Charlie Feigin [28]. This consists of 59 vertebrate species present in the publicly available UCSC 60-way WGA of vertebrates (https://hgdownload.soe.ucsc.edu/goldenPath/mm10/multiz60way/) [37]. The genome of the domestic dog *Canis lupus familaris* was removed from this WGA while genomes of the thylacine *(Thylacinus cynocephalus*), and gray wolf *(Canis lupus)* were added.

We downloaded mouse CDS from Ensembl (version 102, released 2020). With GffRead (version 0.12.7) [49], we retained the longest complete CDS for each protein-coding sequence (parameters -C -Q -K -M) and subsequently extracted fourfold degenerate sites with the msa_view utility from PHAST (version 1.5) [50]. phyloFIT (from PHAST, version 1.5) was then used to estimate the neutral model [50].

We ran phastCons with PHAST (version 1.5) [50], using the –most-conserved option to estimate discrete vertebrate conserved elements from the 61-way alignment, using parameters from the phastCONs track in the UCSC Genome Browser (–expected-length = 45 bp, –target-coverage = 0.30, –rho = 0.30). In determining lineage-specific acceleration of conserved elements, the target lineage(s) is typically masked from the WGA when identifying conserved elements, with the mafSpeciesSubset utility from the UCSC Genome Browser (https://github.com/ucscGenomeBrowser/kent). Masking prevents acceleration in the target lineage from confounding estimates of conservation in the overall alignment. Thus we first masked the genome for each target species (thylacine, grey wolf, Tasmanian devil, giant panda, tammar wallaby and domestic cat) individually from the WGA. We obtained vertebrate conserved elements from this masked WGA, which we then tested for acceleration in the target species. Following these steps, we had distinct sets of conserved elements for each target lineage. Table 2 compares the amount of conserved nucleotide sequence that was tested for acceleration in all target lineages. We determined 75.3 Mb (greater than 96% in each species) of predicted conserved sequence was overlapping across across all six species.

**Table 2.**
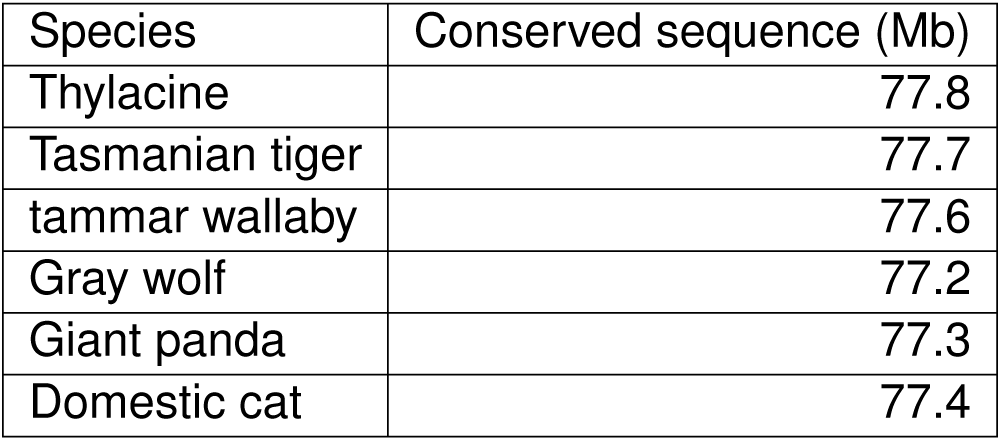
Amount of vertebrate conserved sequence tested for acceleration in each lineage. Prior to estimate of conserved sequence, the target lineage is masked from the WGA.

| Species | Conserved sequence (Mb) |
| --- | --- |
| Thylacine | 77.8 |
| Tasmanian tiger | 77.7 |
| tammar wallaby | 77.6 |
| Gray wolf | 77.2 |
| Giant panda | 77.3 |
| Domestic cat | 77.4 |

We used phyloP (from PHAST, version 1.5) [50] to test for acceleration (mode -ACC, method -LRT. Prior to this, we merged vertebrate conserved within 10 bp of each and filtered out vertebrate conserved elements shorter than 50 bp. We also discarded elements that overlapped with RepeatMasker annotations for the reference mouse mm10 genome (http://www.repeatmasker.org). We applied a Benjamini-Hochberg multiple testing correction with a FDR of 10%.

To overlap different datasets of accelerated regions, we used BEDTools intersect or multiinter (version 2.3.1) [51].

### Defining a test dataset of vertebrate conserved elements for Lift&Add

We downloaded the WGA of 60 vertebrate species as well as the associated neutral model of evolution from the UCSC Genome Browser (https://hgdownload.soe.ucsc.edu/ goldenPath/mm10/multiz60way/) [37]. Alignments were filtered to remove sub-alignments with fewer than 30 species. As above, discrete conserved elements were predicted with the phastCons (PHAST version 1.5) –most-conserved option [2, 50]. BEDtools (version 2.31.1) –merge and –subtract were used to merge elements within 10 bp of each other [51]. We filtered out elements shorter than 50 bp and those overlapping RepeatMasker annotations for mouse mm10 genome. The sequences were labelled with mouse coordinates (chr:start-stop) for identification. The UCSC mafFrags utility (https://github.com/ucscGenomeBrowser/ kent) was subsequently used to extract discrete multiple species alignments (MSAs) for each vertebrate conserved element. The output Multiple Alignment Format (MAF) file was mapped to FASTA with the Galaxy MAF to FASTA tool (https://usegalaxy.org.au/root?tool_id=MAF_To_Fasta1), and split into a separate FASTA file for each vertebrate conserved element.

### Workflow

We constructed a workflow consisting of a series of Snakemake and bash scripts to add sequences from target genomes to existing alignments of conserved regions (Table 3). The workflow is parallelized such that it can be implemented for multiple target genomes and chromosomes at the same time. The scripts and associated configuration files are available at https://gitlab.unimelb.edu.au/igr-lab/thylacine_ canid_convergence_reanalysis/-/tree/master/code/Lift_and_Add. Details for each step are given in the following sections.

**Table 3.** Table 6: Summary of scripts used to add genomic sequences from new genomes to alignments of vertebrate conserved elements.

| Scripts | Steps |
| --- | --- |
| 1.make_dir.sh | Makes individual output directories for each target species, per chromosome. |
| 2.Snakefile | Maps with liftOver (v.1.6.3) and then Liftoff(v.0.1.6) elements from reference to target genome. Extracts sequences of target coordinates from their genomes with BEDtools (v.2.31.1) and splits the output into individual fasta files per conserved element with UCSC faSplit. |
| 3.mv_files.sh | Moves individual conserved element fasta files into the appropriate directory. Also changes the header of each fasta file to the correct species name. |
| 4.combine_seqs.sh | Combines all mapped sequences (across different species) for each conserved element into a single fasta file. |
| 5.mafft_add.sh | Adds new genome sequences to existing alignments with MAFFT. |

#### Snakefile, key steps

- **Mapping conserved elements from mouse to opossum, Tasmanian devil, tammar wallaby** We used UCSC liftOver (version 1.6.3) to map coordinates of the vertebrate conserved elements from the mouse (mm10) genome to the tammar wallaby (macEug2), opossum (monDom5), Tasmanian devil (sarHar1) and domestic dog (canFam3) genomes. All chain files were downloaded from the UCSC Genome Browser (https://hgdownload.soe.ucsc.edu/goldenPath/mm10/liftOver/). We set the liftOver parameter -minMatch at 0.5, which requires 50% coverage for mapped sequences between the reference and target genomes. This was set at a more conservative threshold than the default 0.10 for inter-species coordinate conversion with liftOver, in order to obtain a high confidence test dataset. The output BED files are mapped to a Gene Transfer Format (GTF) with bed2gff.py (from CGAT [52], https://github.com/CGATOxford/cgat/blob/master/ CGAT/scripts/bed2gff.py).
- **Mapping conserved elements from Tasmanian devil/tammar wallaby to new marsupial genomes with Liftoff** We next used two rounds of Liftoff [34] to map coordinates of vertebrate conserved elements from a reference species to a closely related target species. Liftoff uses local alignment with minimap2 [47] to map sequences between the query and target genome. Inputs required include the query and target genomes and the GTF file of elements to be mapped. The Liftoff output consists of a GTF file of mapped elements and a list of unmapped ones. We used the -exclude-partial parameter to exclude partial alignments from the output as recommended by Liftoff—these are alignments that have lower than 50% alignment coverage and/or sequence identity between the query and target. We performed two rounds of Liftoff—in the first round, we mapped elements with the parameter -flank set at 0 (default). This parameter adjusts the amount of flanking genomic sequence added to a query element as a proportion of its total length. Next, for unmapped elements, we repeat Liftoff with -flank set a 1, which adds the maximum possible amount of flanking genomic sequence to the query element. We combined the mapped elements from both rounds of Liftoff into a single GTF file.
- **Getting sequences from each target genome.** We converted the combined GTF file of mapped elements from Liftoff to BED with BEDOPS (version 2.4.4.1) gtf2bed [53]. We then extracted the nucleotide sequences for all the mapped elements with the BEDTools getfasta utility [51]. We obtained individual FASTA files per element with the UCSC faSplit utility (https://github.com/ucscGenomeBrowser/kent). Lastly, an output table is created summarising the species to which each element was mapped.

#### Adding new sequences to multiple species alignments of vertebrate conserved elements with MAFFT

For each element that was successfully mapped to at least one species, we generated compiled FASTA files consisting of sequences from all mapped species (3.mv files.sh, 4.combine_seqs.sh, Table 3). We then added the new species sequences to the original 60-way alignments with MAFFT –add (5.mafft_add.sh, 3). We used the following parameters:

- –adjustdirectionaccurately, to account for reverse complement sequences.
- –keeplength, to retain the structure of the original MSA, such that only conserved sequences are kept and any flanking sequence that is not conserved is trimmed.
- –localpair, L-INS-i local pairwise alignment strategy, the most accurate MAFFT algorithm appropriate for alignment of fewer than 200 sequences.

As quality control for this step, we computed pairwise distances between species alignments with the ape package (version 5.8-1) in R (version 4.40), using model = "raw" in the function dist.dna(). Under this model, pairwise distance is simply defined here as proportion of sites that differ between a pair of aligned sequences. For every alignment of a conserved element, we calculated the pairwise alignment distance between Tasmanian devil (sarHar1) or tammar wallaby (macEug2) and each new marsupial species in the alignment. For comparison, we also calculated the pairwise alignment distances between the Tasmanian devil (sarHar1), tammar wallaby (macEug2) and opossum (monDom5).

### Using RERConverge to correlate evolutionary rates shifts with thylacine-wolf convergence

We used the following RERconverge (version 0.1.0) walkthroughs—1) https://github.com/ nclark-lab/RERconverge/blob/master/vignettes/PhangornTreeBuildingWalkthrough.pdf and 2) https://github.com/nclark-lab/RERconverge/blob/master/vignettes/ FullWalkthroughUTD.pdf. Firstly, we estimated for each MSA of a vertebrate conserved element a phylogenetic tree with the estimatePhangronTreeAll() function. We chose the widely-used general time reversible (GTR) substitution model for phylogenetic estimation, which is also recommended for DNA sequences by RERConverge. RERConverge then was used to estimate the consensus tree from the individual trees, following by calculation of the relative evolutionary rates (RERs) for all branches for each genomic element withgetAllResiduals(). The old genomes for the Tasmanian devil (sarHar1), wallaby (macEug2) and domestic dog (canFam3) were excluded when calculating these RERs to remove redundancy, as newer genomes of these species (or of a closely-related subspecies for the dog) were also present in the alignments.

We encoded thylacine-wolf convergence as a binary phenotype with the function foreground2tree(), setting clade = ‘terminal’, as we were testing for rate shifts only in the terminal thylacine and wolf branches and not in the internal branches leading to the thylacine-wolf common ancestor [42]. Supplementary Figure 10 displays the tree model for the binary phenotype, with the thylacine and wolf foreground lineages highlighted. With the function correlateWithBinaryPhenotype(), the RERs were associated with the binary trait. We set the parameter min.pos = 20, which requires a minimum of 20 species to be present in a MSA included in this analysis. The default setting is min.pos = 10, but given that the maximum number of species in my analysis was 73, this was increased to 20 to exclude highly sparse alignments.

### Plots and statistical analysis

All plots and statistical testing detailed were implemented in R (version 4.2.1). Analysis scripts are available at https://gitlab.unimelb.edu.au/igr-lab/thylacine_canid_convergence_ reanalysis/-/tree/master/analysis.

## Acknowledgements

We thank Charlie Feigin, Laura E. Cook and members of the Gallego Romero Lab for their analysis advice. NS was supported by a Melbourne Research Scholarship and the David Lachlan Hay Memorial Grant from the University of Melbourne. St Vincent’s Institute acknowledges the infrastructure support it receives from the National Health and Medical Research Council Independent Research Institutes Infrastructure Support Program and from the Victorian Government through its Operational Infrastructure Support Program.

## Author contributions

- Conceptualization: NS, IGR
- Formal analysis: NS
- Investigation: NS, IGR
- Methodology: NS
- Resources: IGR
- Supervision: IGR
- Visualization: NS
- Writing: NS, IGR

## Supplementary Figures

**Supplementary Figure 1.**
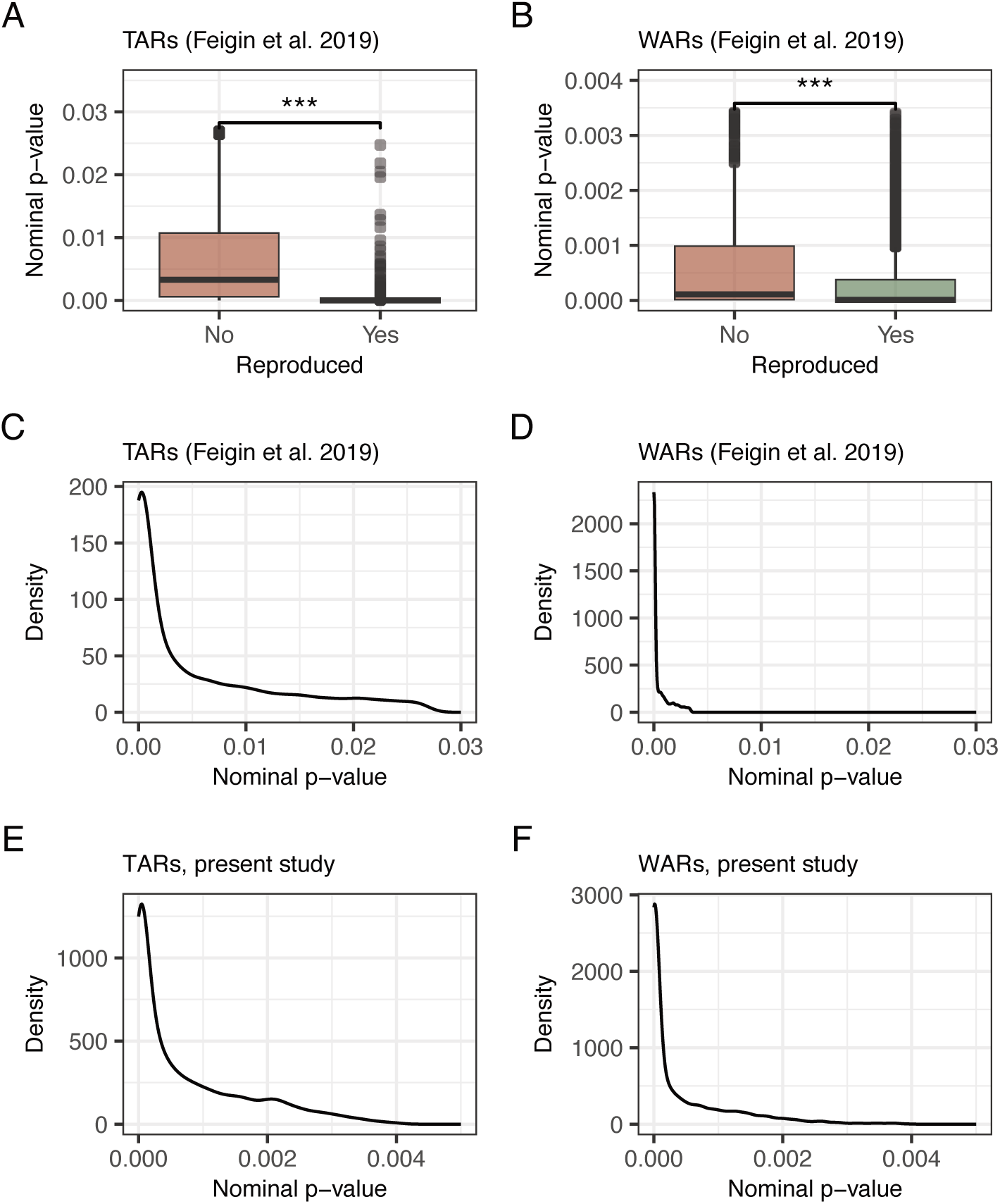
*P*-values for significant acceleration. **A.** Differences in nominal *p*-values distributions between TARs from Feigin et al. [28] reproduced and not reproduced in our analysis **B.** Differences in nominal *p*-values distributions between WARs reproduced and not reproduced in our analysis. **C.** Overall nominal p-value distributions for TARs, 2019, **D.** WARs, 2019, **E.** TARs, 2025, and **F.** WARs, 2025. The TARs had significantly larger nominal *p-*values than the WARs in both the 2019 study and the present 2025 study (pairwise t-test *p-*value = = *<* 2.2 *×* 10*^−^*^16^).

**Supplementary Figure 2.**
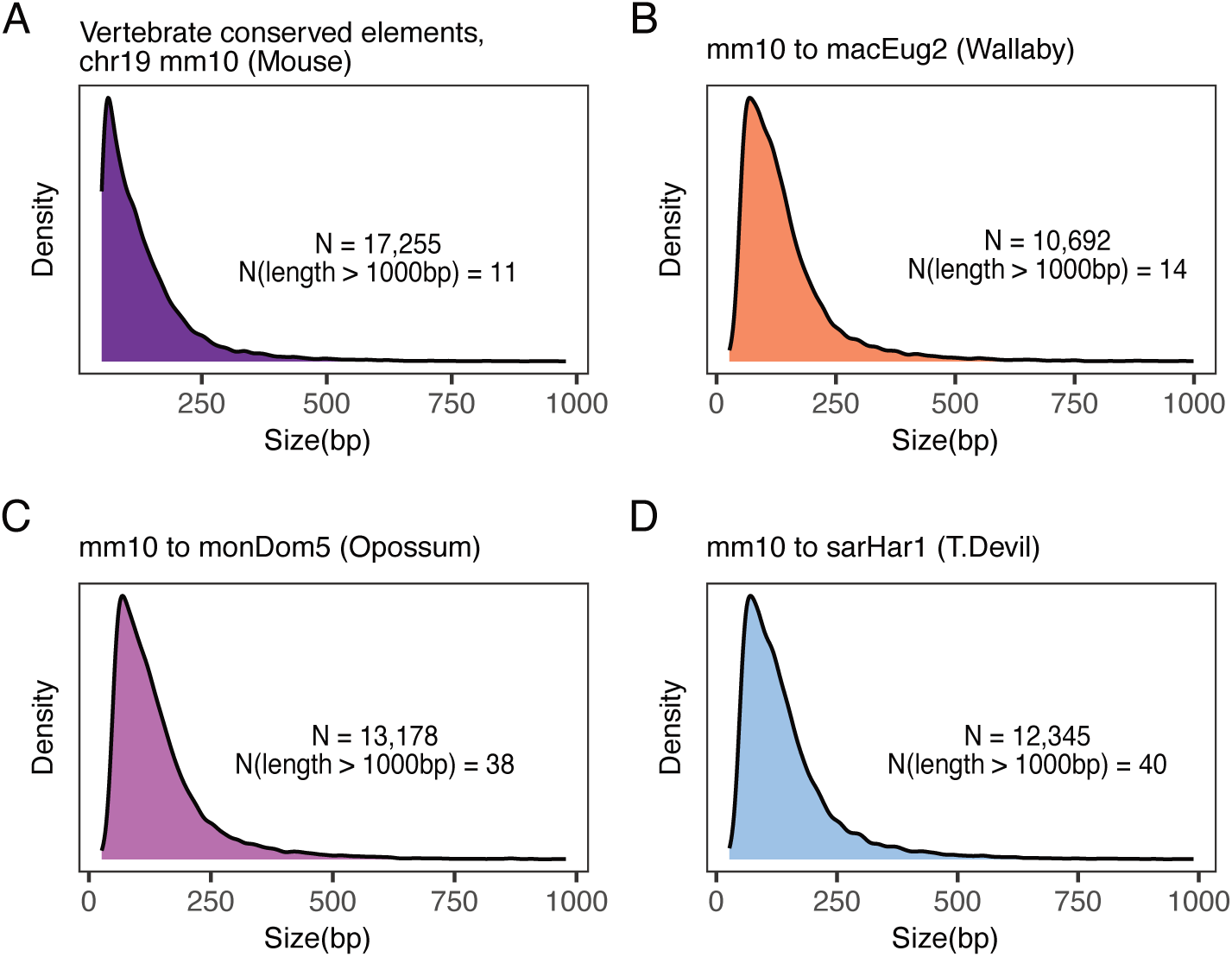
Size(bp) of vertebrate conserved elements. **A.** as output by PHAST in mm10 mouse coordinates and after liftOver to the **B.** wallaby macEug2 genome **C.** opossum monDom5 genome and C. Tasmanian devil sarHar1 genome.

**Supplementary Figure 3.**
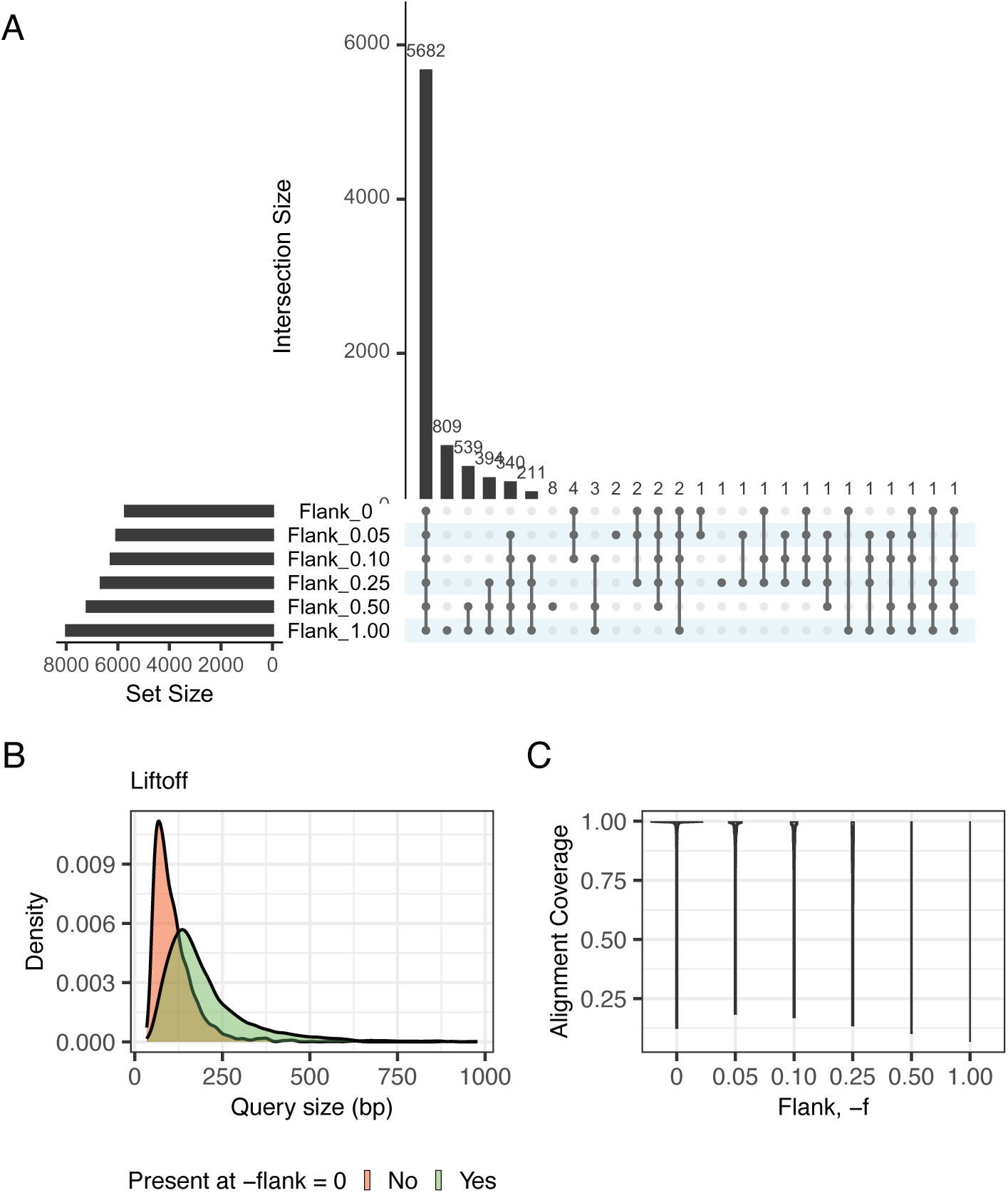
**A.** Intersection of Liftoff output at different thresholds of -flank (0, 0.05, 0.10, 0.25, 0.50 and 1). **B.** Distribution of sizes for opossum query elements that are mapped to devil at -flank = 0 or -flank *>* 0. **C.** Distribution of alignment coverage for Liftoff output elements at different threshold of -flank.

**Supplementary Figure 4.**
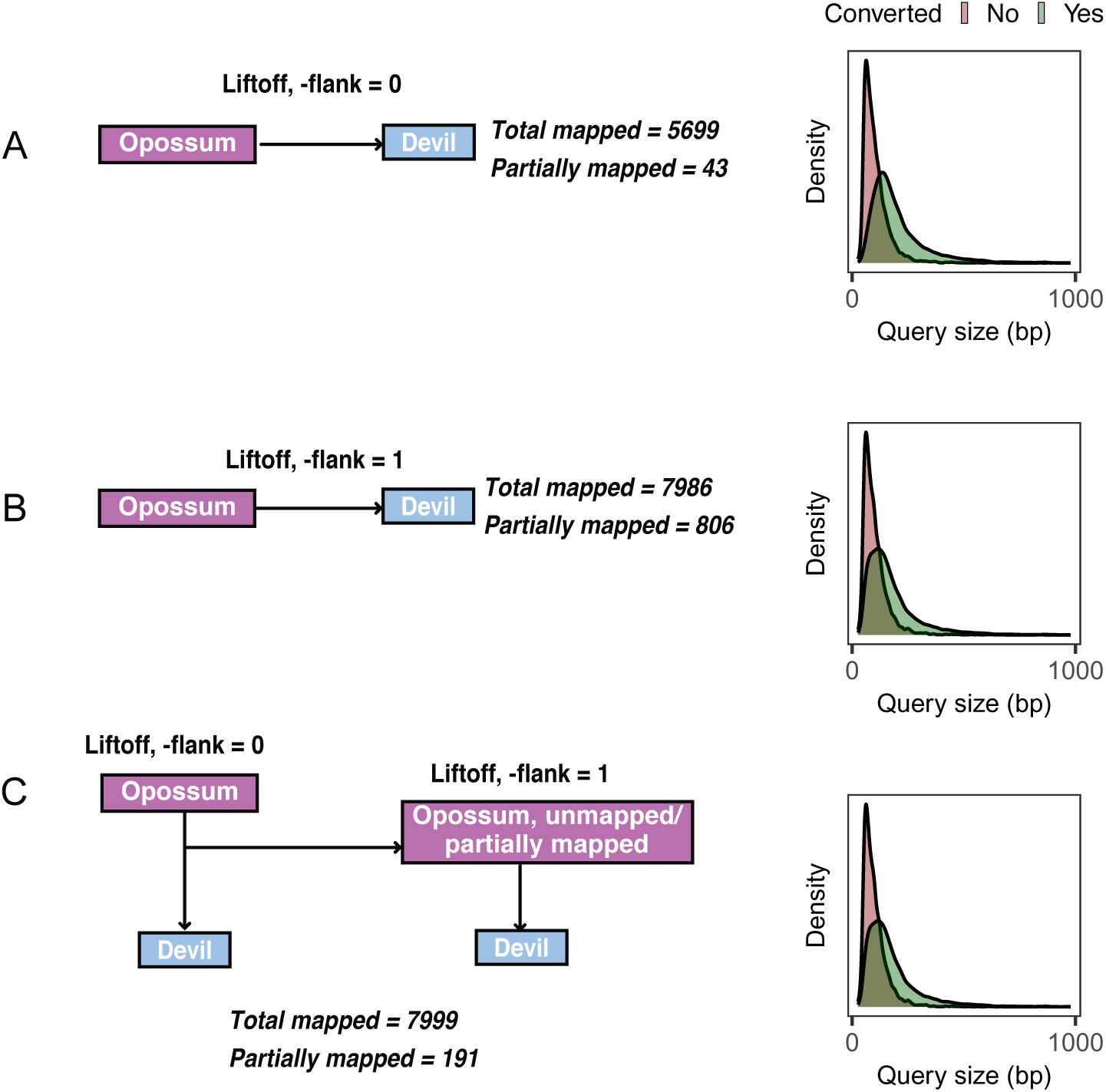
Two-rounds of Liftoff to increase recovery and reduce partial mapping. Summary of Liftoff output with **A.** No flanking genomic sequence added (-flank = 0), **B.** Maximum amount of flanking genomic sequence added (-flank = 1) and **C.** -flank = 0, followed by a second Liftoff for unmapped elements with -flank = 1.

**Supplementary Figure 5.**
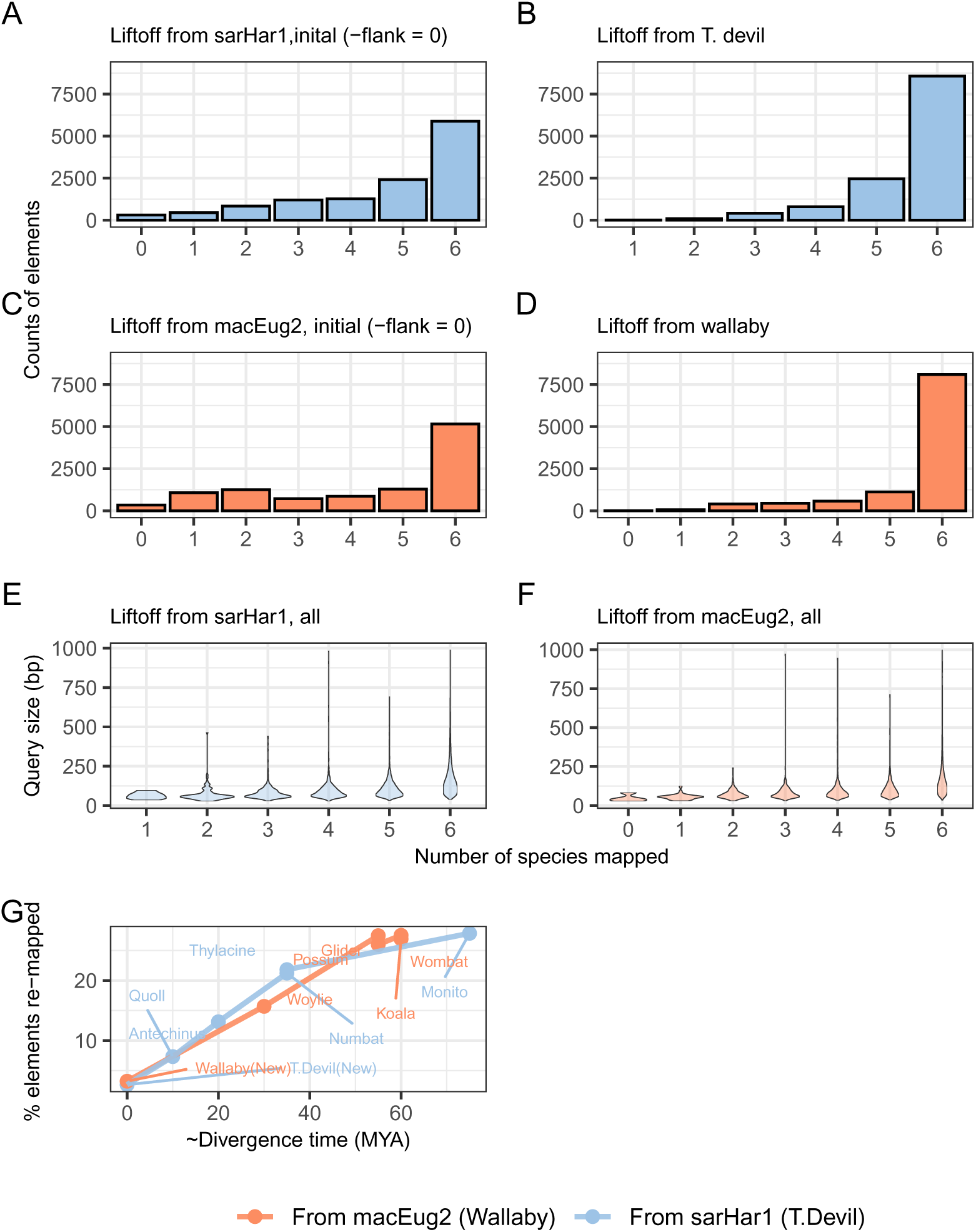
Coordinate conversion from devil (sarHar1) or wallaby (macEug2) to target marsupial genomes. **A and C.** Counts of query elements that were mapped to 0 or more species, with initial Liftoff (no flanking genomic sequence). **B and D.** Counts of query element that were mapped to 0 or more species, after both rounds of Liftoff. **E and F.** Size distribution for query elements that mapped to 0 or more species. **G.** Correlation between estimated divergence between query and target and the percentage of elements mapping only with the addition of flanking sequence (-flank =1).

**Supplementary Figure 6.**
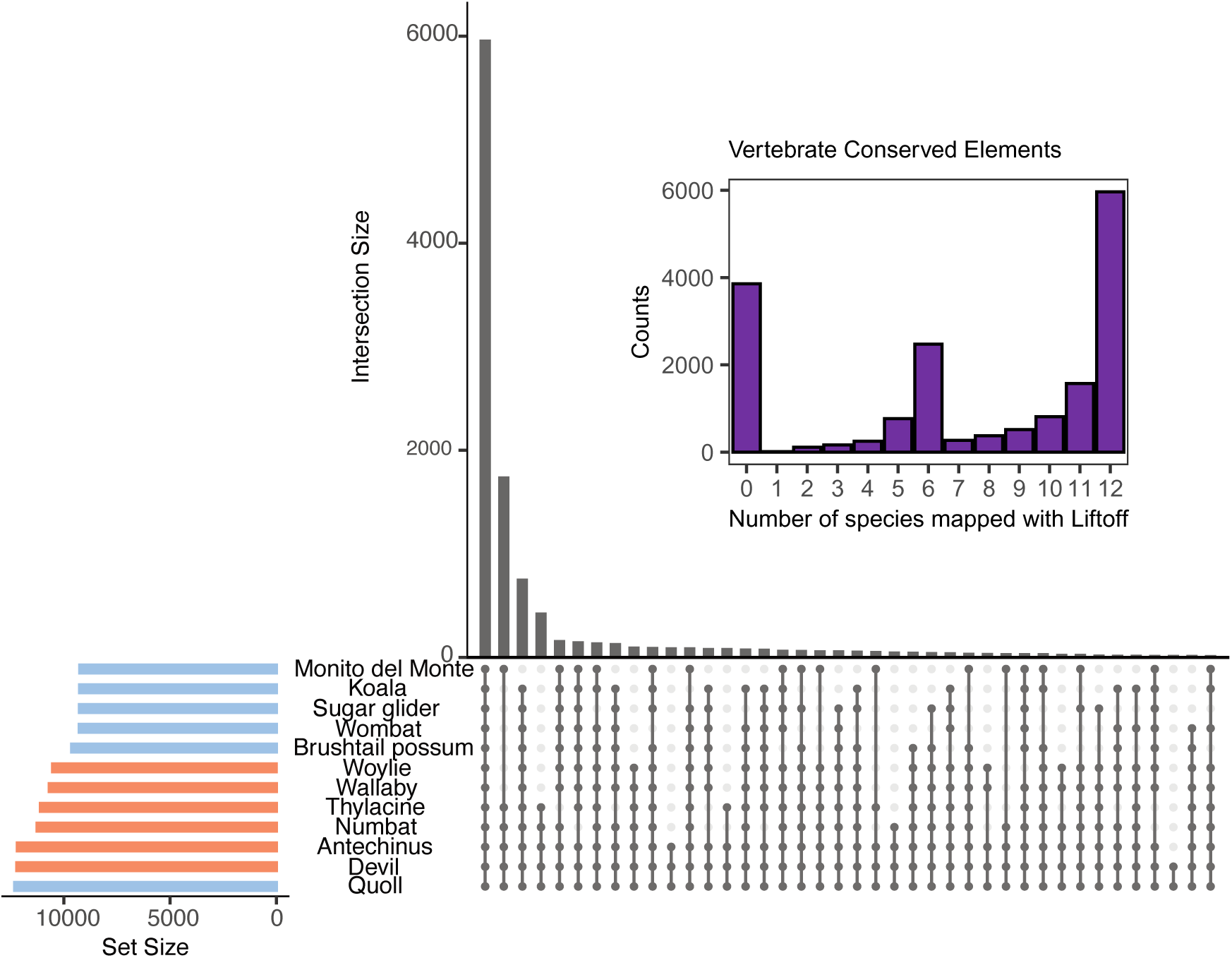
Intersection of Liftoff output across the 12 marsupial target genomes. The histogram displays the number of the original 17,255 vertebrate conserved elements that were mapped to 0 or more species.

**Supplementary Figure 7.**
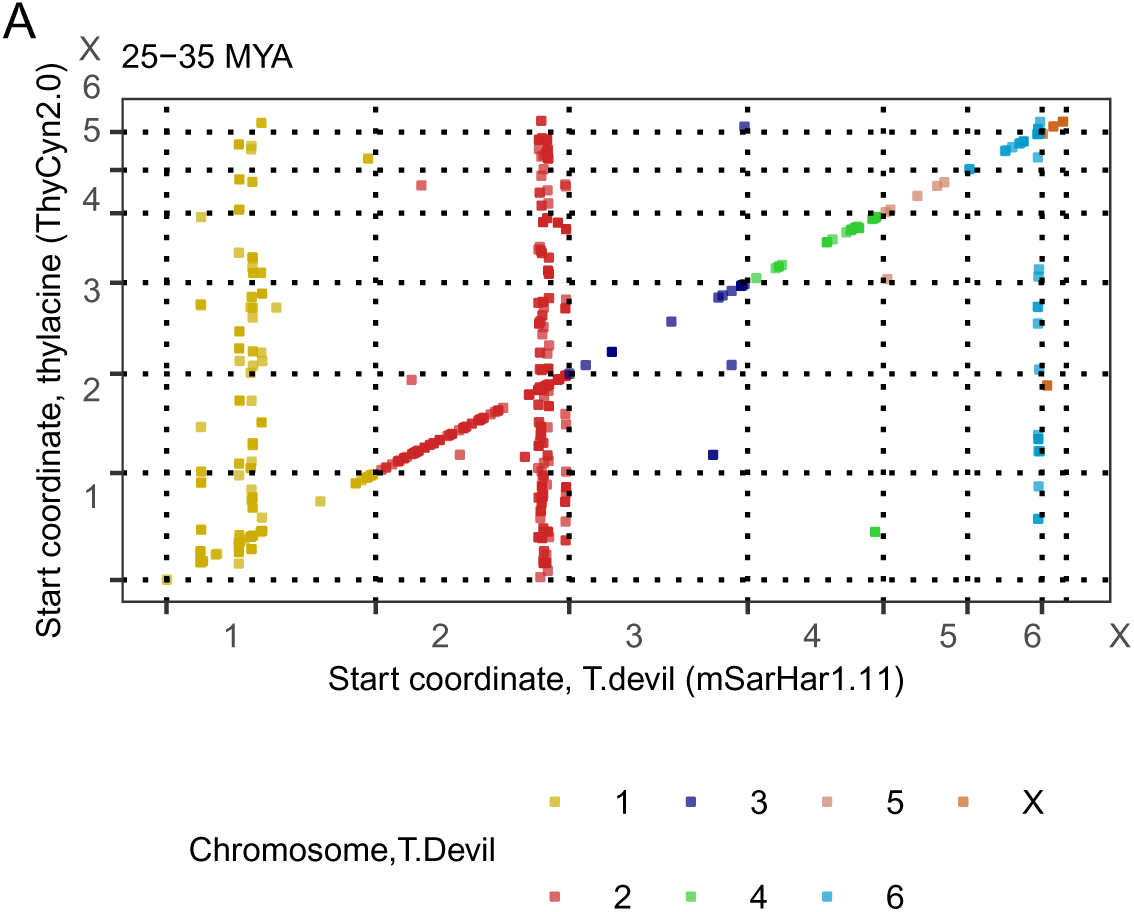
Correlation in genomic coordinates of mapped elements between the Tasmanian devil (mSarHar1.11) and Thylacine (ThyCyn2.0), highlighting regions at which conserved elements are displaying variable mapping between these genomes.

**Supplementary Figure 8.**
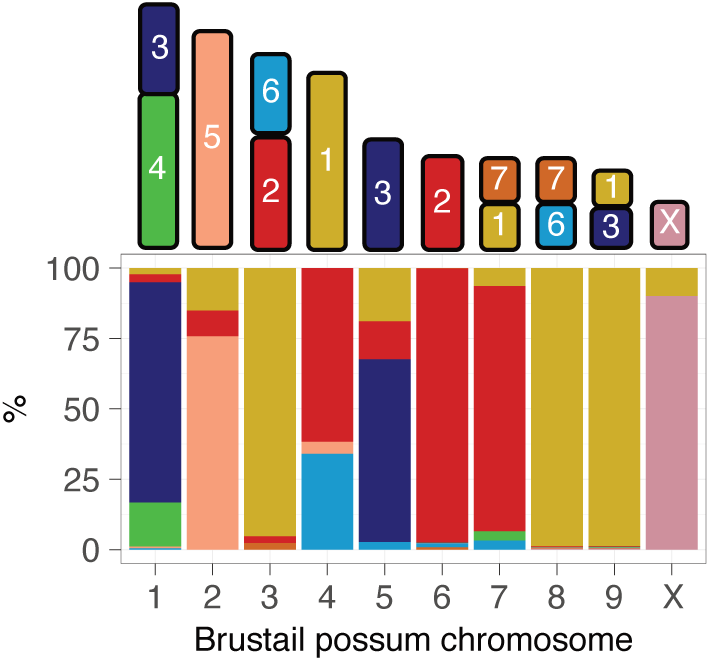
Relationship between the tammar wallaby and brushtail possum karyotypes, as determined by [40], and proportion of test elements mapping to each brushtail possum chromosome, coloured by their location in the tammar wallaby genome.

**Supplementary Figure 9.**
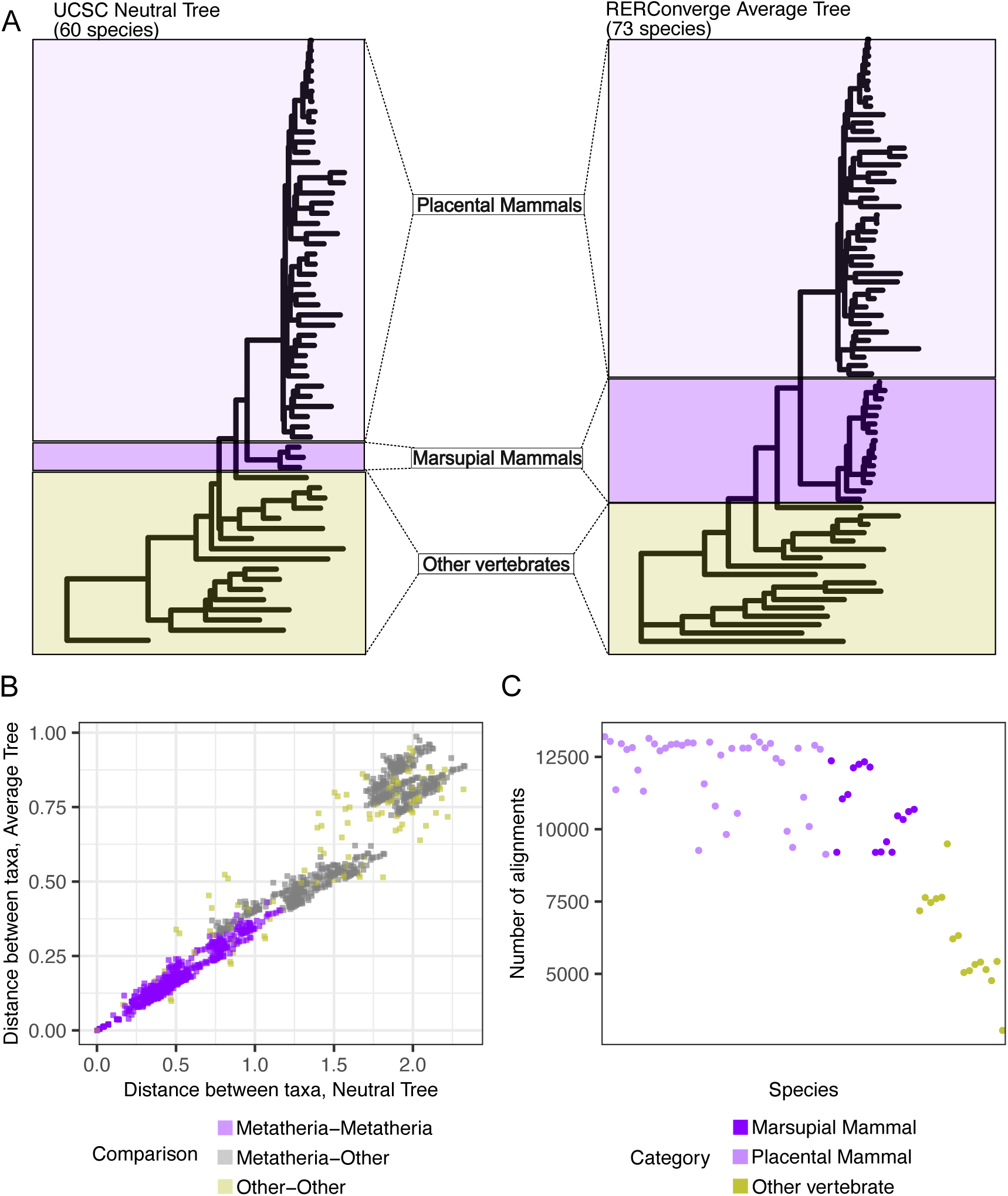
Average tree, RERConverge. **A.** Neutral model of evolution for the 60 species WGA (left) and consensus tree estimated from alignments of 13,220 vertebrate conserved elements by RERConverge (right). **B.** Distances between terminal nodes in the neutral tree versus those in the average tree. **C.** Number of alignments in which each species is present.

**Supplementary Figure 10.**
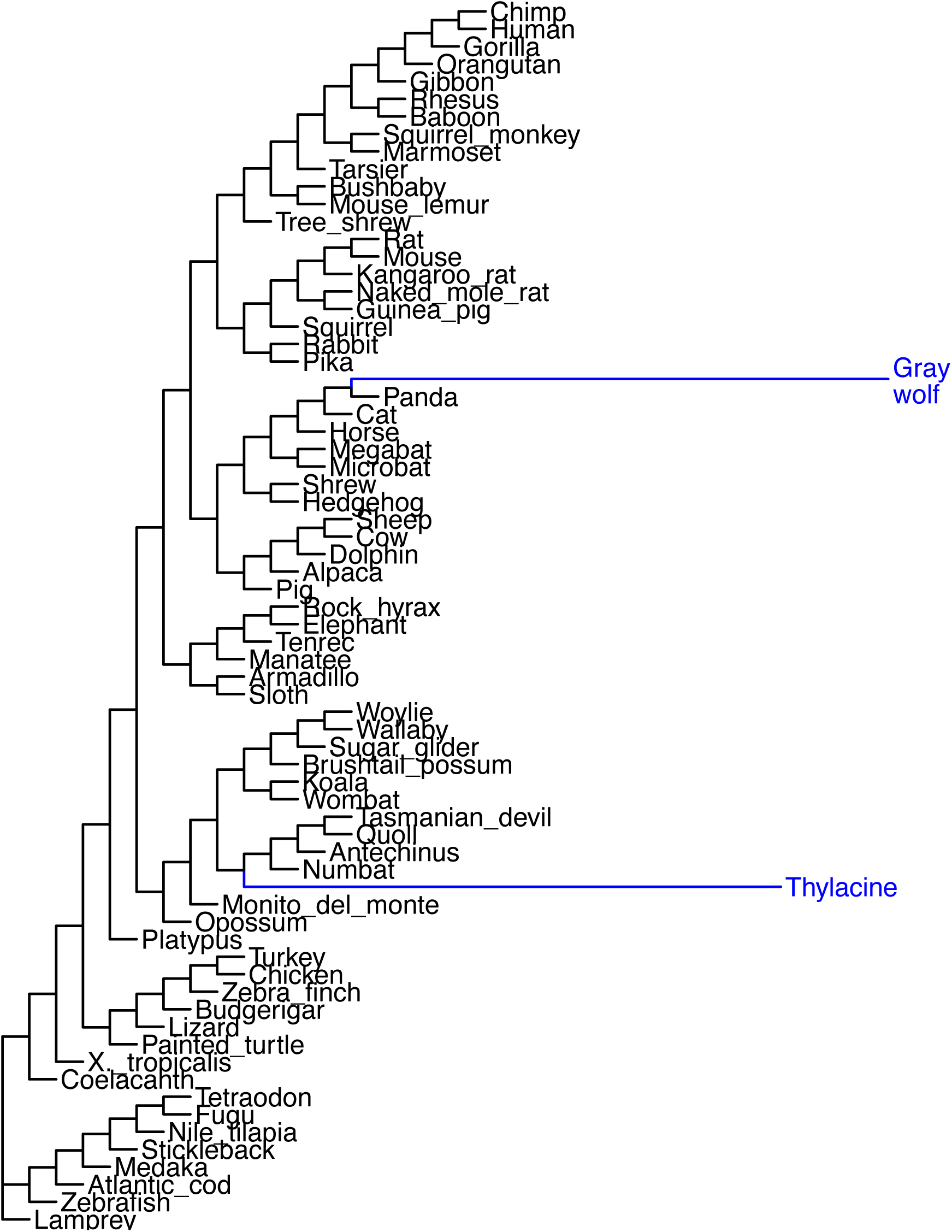
Binary trait tree. Used as input in the RERConverge trait association analysis. In this tree, the foreground convergent lineages have branch lengths of 1 while the background lineages have branch lengths of 0.

## Supplementary Tables

**Supplementary Table 1**: Number of significantly accelerated regions determined in both the Feigin et al. [28] and the present study.

**Supplementary Table 2**: List of query genomes for Liftoff.

**Supplementary Table 3**: liftOver and Liftoff of vertebrate conserved elements from the gray short-tailed opossum genome (monDom5) to the Tasmanian devil genome (sarHar1).

**Supplementary Table 4**: Liftoff output at different threshold of -flank.

**Supplementary Table 5**: List of target marsupial assemblies.

**Supplementary Table 6**: Number of test elements from Wallaby chromosomes that mapped to each Brushtail possum chromosome with Liftoff.

**Supplementary Table 7**: Pairwise comparisons of alignment distances to the Tasmanian devil (sarHar1).

**Supplementary Table 8**: Pairwise comparisons of alignment distances to the Tammar wallaby (macEug2).

